# CWLP and PRP940 form plasma-membrane nanodomain complexes with aquaporins, interact with PP2A and contribute to dehydration tolerance

**DOI:** 10.1101/2022.10.20.512913

**Authors:** Arik Honig, Haviva Eilenberg, Ellen Zuther, Dror Avisar, Oran Erster, Gad Eshkol, Hanan Stein, Noam Reznik, Silvia Schuster, Eduard Belausov, Muhammad Abu-Abied, Takashi Hotta, Yuh-Ru Julie Lee, Bo Liu, Einat Sadot, Csaba Koncz, Aviah Zilberstein, Yosef Fichman

## Abstract

The C-type hybrid-proline-rich protein (HyPRP) AtCWLP and its homolog AtPRP940 are referred as cell wall (CW)-plasma-membrane (PM) linker proteins, but little is known about their functions. Here we show that N-terminal proline-rich domains of CWLP and PRP940, containing glycosylated hydroxyproline residues, contact the CW, while their C-terminal 8CM domains function as PM-scaffolds. Both proteins are detected in PM nanodomains (PM-ND) and show co-localization and co-immunoprecipitation with aquaporins PIP2;1 and PIP2;7. Inhibition of actin polymerization by latrunculin B promotes CWLP-endosome appearance, while blocking the actomyosin-based transport by a truncated form of myosin XI-K relaxes lateral boundaries of CWLP-PIP2;1 PD-NDs. Mass spectrometry data indicate that CWLP co-purifies with dynamins implicated in fission of endocytic PD-ND invaginations. Lack of co-localization and co-immunoprecipitation with aquaporin-binding flotillin (FLOT2) indicates that CWLP and PRP940 mark a new distinct type of PM-ND. Yeast two-hybrid and co-immunoprecipitation assays demonstrate that CWLP and PRP940 interact with multiple aquaporins and several protein phosphatase PP2A-B’’ regulatory subunits. By preventing irreversible separation of CW and PM, and likely assisting PP2A-mediated dephosphorylation of aquaporins and closure of their water channels, overexpression of CWLP confers tolerance to plasmolysis, dehydration and freezing in Arabidopsis and to water shortage in potato plants.

**Summary Statement:** Arabidopsis Hybrid-Proline-Rich Proteins CWLP and PRP940 occur in association with dynamins, recruit PP2A protein phosphatases to aquaporin water channels in plasma-membrane (PM) nanodomains and elevate tolerance to cellular dehydration.

## 1 INTRODUCTION

Hybrid Proline-Rich Proteins (HyPRPs) unique to gymnosperms and angiosperms comprise a family of highly variable proteins with diverse cellular localizations. Arabidopsis CWLP (At2g22120; PRP16) and its homolog PRP940 (At3g10940; PRP15) are classified into the C-type family of HyPRPs (1, 2). HyPRPs are also considered as the third group of large HRGP (Hydroxyproline-Rich Glyco-Protein) family, including extensins and arabinogalactan proteins [AGPs; (3, 4)], all carry proline-rich domains (PRDs) with a great variety of short repeats (5). These repeats contain hydroxyproline (O, or HyP) residues resulting from Pro hydroxylation and O-glycosylation by successive activities of prolyl 4-hydroxylases and glycosyltransferases in the endoplasmic reticulum (ER) and Golgi (6–8). HRGPs possessing N-terminal secretory signal peptides (SSPs) traffic through the ER-Golgi-trans-Golgi Network (TGM), and their delivery to the PM by either clathrin-coated or secretory vesicles is assisted by the actomyosin network in the Conventional Secretory Pathway (CSP). Cell wall proline-rich proteins (CW-PRPs) lacking SSPs and some HGRPs under stress are delivered to the CW and PM or extracellular space by Unconventional Secretory Pathways [USPs; (9, 10)]. Through their glycosyl moieties, HRGPs and CW-PRPs interact with diverse CW components (11, 12), assisting CW assembly and remodeling (13). HRGPs and CW-PRPs form various complexes with other PM-proteins, e.g., aquaporins, membrane transporters, ATPases, receptor protein kinases, and diverse signaling factors, and thus play pivotal roles in cellular responses to hormonal, biotic, and abiotic stress stimuli. These signaling complexes form distinct PM clusters called micro/nanodomains (PM-NDs), which are particularly rich in sterols and phyto-sphingolipids (Glycosyl Inositol Phospho Ceramides, GIPCs), as well as S-acylated, glycosylated and poly-phosphoinositide-binding proteins (14–17). These nanodomains form detergent-resistant membrane entities (DRMs or lipid rafts), and restrict lateral diffusion of PM resident proteins (18–20). PM-NDs are characterized by a typical collection of DRM marker proteins (21–23). Cell wall contacts of DRM resident proteins appear to play an important role in determining the size and positioning of PM-NDs. Replacement of CW contact domains with GFP relaxes lateral mobility and extends the boundaries of several DRM-resident “linker/scaffold” proteins (24–26). Hormonal and stress signals stimulate cytoplasmic invagination of PM-NDs forming characteristic PM-pits, which are internalized to clathrin-coated or secretory endocytic vesicles, and recycled to the PM respectively by predominant clathrin-mediated (CME) and stress-induced clathrin-independent (CIE) endocytosis pathways (10, 27–29). Like in mammals, endocytosis in plants is aided by cortical actin filaments (AFs). In plants, inhibition of actin polymerization does not prevent dynamin-mediated scission of endocytic membrane pits. In plant CME pathway, actin nucleation is driven by the Arp2/3 complex (30, 31), whereas in CIE, this is likely dependent on dynamins/ Dynamin-Related (DRP) GTPases (32–35). Plant CIEs appear to be less sensitive to latrunculin B (LatB)-mediated inhibition of actin-polymerization, probably due to different properties of actin- and dynamin-interacting PM-ND scaffold proteins. For instance, the abundant PM-ND-resident remorin proteins (e.g., REMORIN1;2 and 1;3) interact with CW-galacturonides through their N-terminal proline-rich domains, whereas via their C-terminal transmembrane domains bind to formins (e.g., FORMIN FH6), promoting actin nucleation (36). PM-NDs marked by remorins recruit numerous receptor and transporter proteins, including e.g., the auxin transporter PIN2 and flg22 receptor FLS (37–39). Endocytosis of PIN2 and FLS is notably reduced by LatB, but not abolished completely, although LatB prevents their endocytic trafficking and PM recycling (39, 40). LatB seems to have a weaker effect on dynamin-dependent endocytosis of FLOT2 (FLOTILLIN 2; another nanodomain marker and binding partner of FLOT1 and aquaporins in Arabidopsis), and its mammalian homologs, which interact with cortical F-actin, dynamins and a range of regulatory proteins (25, 41–46). Flotillins also play a key role in ER to PM-ND trafficking and endocytosis of nonspecific Lipid Transfer Proteins (nsLTPs) that are linked to the PM by glycosylphosphatidylinositol (GPI)-anchors (47–49).

All 28 Arabidopsis HyPRPs including CWLP and PRP940 consist of N-terminal SSPs, PRDs and C-terminal hydrophobic domains with highly conserved motifs of eight Cys residues (8CM). Compared to extensins and AGPs, HyPRPs contain fewer hydroxyproline residues and thus have lower O-glycosylation levels (5, 50). The 8CM domain is not unique to HyPRPs; it is present in more than 500 plant proteins, including different types of nsLTPs, prolamins, amylases, trypsin inhibitors, and albumin 2S-like seed storage proteins (29, 51–53). In sequenced plant genomes, *HyPRP* genes are found in chromosomal clusters. Due to the sequence heterogeneity of their PRDs, classification of Arabidopsis HyPRPs into seven subgroups is primarily based on their chromosomal locations and similarities among their 8CM domains (1). Six out of seven genes in Arabidopsis *EARLI1* subfamily of HyPRPs are thus clustered in chromosome IV. Ectopic expression of *EARLI1* (At4g12480) is reported to elevate freezing tolerance in yeast and plants (54), while AZI1 (AZELAIC ACID INDUCED 1) encoded by adjacent *EARLI1* paralog (At4g12470) is found to regulate salt tolerance in interaction with MPK3 (MITOGEN-ACTIVATED PROTEIN KINASE 3); (55). Moreover, AZI1 and EARLI1 are shown to interact with nsLTP DIR1, and mediate the transport of SAR (Systemic Acquired Resistance)-inducing azelaic acid and its precursors (56–58). Recently, AZI1 and other small AZI1-LIKE (AZL) HyPRPs (i.e., carrying less than 180 amino acids) were reported to occur in ring-like structures around the chloroplasts, and detected in chloroplast membrane fractions by mass spectrometry. AZI1 and AZLs are proposed to function in interaction with nsLTPs (59) as tethering factors that define Membrane Contact Sites (MCSs) for bidirectional non-vesicular lipid exchange through stromule-like structures between the ER and chloroplasts (60–64).

In contrast to the small AZL proteins, the size of C-type HyPRPs, including CWLP/PRP16, PRP940/PRP15, PRP14 (At1g62500), At4g15160, and potentially also At3g22142 and At4g22485 (1) exceeds 275 amino acids due to longer lengths of their PRDs. When coining the term “cell wall-membrane linker” protein for *Brassica napus* PROLINE-RICH PROTEIN (BnPRP), the founding member of C-type HyPRPs and the closest homolog of Arabidopsis CWLP and PRP940, Goodwin et al., (65) predicted that BnPRP is anchored to the cell wall by glycosylation of PRD residues, while its hydrophobic C-terminus embedded in the PM. This prediction was supported by bioinformatics analysis of plant HPRGs (66). AtCWLP was first identified as a putative PM protein by a yeast-based screening for secretory signal peptides using a Arabidopsis cDNA library (67). Subsequent mass spectrometry studies detected both CWLP and PRP940 in purified CW fractions (50, 68, 69); (http://www.polebio.lrsv.ups-tlse.fr/WallProtDB/). CWLP homologs in other species, such as the cold-induced Brassica BnPRP (65), salt-induced alfalfa MsPRP2 (70), and cold- and arachidonic acid-induced tomato DEA1 (71) have been implicated in the modulation of osmotic stress tolerance. Cellular accumulation of HyPRPs appears to correlate with stress-induced elevation of free proline content (65, 70, 72–74). As proline is one of the major osmolytes accumulating in plants during abiotic and biotic stresses (75–77), it is proposed that the availability of high levels of free Pro might support the synthesis of HyPRPs (74, 78). Nonetheless, the proposed CW-PM linker role of AtCWLP homologs and their exact functions in regulating osmotic stress tolerance remain to be tested.

Here we show that overexpression of AtCWLP and AtPRP940 in Arabidopsis and tobacco enhances tolerance to hyperosmotic stress-induced plasmolysis illustrating their genuine CW-PM linker functions. Furthermore, both proteins interact, co-immunoprecipitate, and co-localize with aquaporins in novel type of PM-nanodomains, which lack aquaporin-recruiting flotillin. CWLP is found in association with dynamin-related proteins and recruits multiple PP2A-B’’ phosphatase regulatory subunits to its complex with aquaporins. Enhanced dehydration tolerance conferred by CWLP and PRP940 overexpression in Arabidopsis and potato and better freezing withstanding in Arabidopsis suggest that these CW-PM linker proteins contribute to the reduction of cellular water loss by promoting the closure of aquaporin water channels through assisting PP2A-mediated dephosphorylation.

## 2. MATERIALS AND METHODS

### 2.1 Construction of CWLP expression cassettes and deletion mutants

Construction of gene expression cassettes CWLP-HA, CWLP-GFP, GFP-CWLP, CWLP-Δ326-334, Δ25-158-CWLP-GFP, PRP940-GFP, PP2A-B”γ-YFP and FLOT2-GFP, all driven by the CaMV 35S promoter, is described in the Supporting Information (SI). Schematic maps of the main constructs are displayed in Fig. S10.

### 2.2 Plant transformation, selection, dehydration and cold stress conditions

Binary vectors carrying the gene constructs were introduced by electroporation into *Agrobacterium tumefaciens* GV3101 (pMP90RK) (79) and transformed into *Arabidopsis* plants by the floral-dip method (80). Transient (co)-expression of proteins was performed according to (81) by infiltration of leaves of 7-8-week-old *Nicotiana benthamiana* plants with *Agrobacterium* GV3101 (pMP90RK) harboring the binary vectors with the different gene constructs. Injected areas were examined after 48 h by confocal microscope or used for Co-IP assays as described in SI.

A homozygous *cwlp* T-DNA knockout mutant was identified in the GABI-KAT collection (82). In the GABI_382D01 *cwlp* mutant, the T-DNA insertion was localized 715 bp downstream of the translation start in the intron-less *CWLP* coding sequence of 1005 bp. Wild type (WT) *Arabidopsis* (Col-0) and transgenic seedlings were germinated on MS medium with 0.8 % agar in the absence or presence of 50 μg/ml kanamycin or 15 μg/ml hygromycin, according to the relevant selection marker. Potato leaf-disc transformation and plant regeneration were performed according to (83). Arabidopsis and potato (*Solanum tuberosum*, cv Desiree) plants were grown in pots under 8 h light (100 μmol.s^−1.^m^−2^) / 16 h dark (SD), or 16 h light (60 μmol.s^−1.^m^−2^)/ 8 h dark (LD) photoperiod at 25°C. Dehydration experiments were performed with well-watered 4-week-old Arabidopsis plants grown in pots containing equal soil weight, using SD conditions and 60 % RH in Percival growth chambers. Aerial plant size and PSII activity were measured daily using a PlantScreen High-Through-Put Phenotyping system (Photon Systems Instruments) as described by (84). Additional dehydration experiments were performed using large soil-containing trays in which WT and homozygous CWLP-GFP plants were grown together for eight weeks under SD conditions at 25°C and then subjected to water withholding for 21 days. Potato dehydration experiments were carried out using five WT and CWLP-GFP plants of similar size grown under LD conditions in pots with equal soil content, which were subjected to water withholding for 21 days. Cold stress was imposed by transferring 4-week-old plants grown in pots under LD conditions at 25°C, to 4°C for 96 h under LD light regime and then transferred back to 25°C for recovery. 42-d-old plants were exposed to freezing temperatures in the range of -1 to -22°C, with or without acclimation for 14 days at 4°C, and then subjected to thawing under controlled conditions. LT_50_ values of detached mature leaves were determined as described (85). Plasmolysis tests were performed with 10-day-old Arabidopsis seedlings submerged in either 30 % glycerol or 0.6 M sorbitol for 20-30 min. Leaves were then mounted on microscope slides in the same solution. Four leaves from each plant line were examined by confocal microscopy, and the experiments were repeated at least three times.

### 2.3 Monitoring gene expression

RNA extraction and northern blot analyses were performed according to (86). As a probe in filter hybridization experiments, *CWLP*-cDNA fragment (100 ng) was labeled by random-priming with [α-^32^P]dCTP (3000 Ci/mmol, 10 mCi/ml; Amersham Pharmacia). Measurement and histological visualization of GUS activity were performed as described (86). For quantitative measurements, RNA extraction, cDNA synthesis, and qRT-PCR reactions were performed according to (84). CWLP and PRP940 PCR primers are listed in Table S2.

### 2.4 Membrane protein fractionation, detection by co-immunoprecipitation and Western blotting, and mass spectrometry of purified CWLP-GFP

Arabidopsis seedlings (15-d-old) were subjected to differential-detergent protein extraction according to (87) with minor modifications described in SI, Section 6. Equal amounts of Triton-X100 insoluble (DRM), Triton-X100 soluble (TSM), and cytosolic supernatant (Sup) protein fractions were resolved by SDS-PAGE and analyzed by immunoblotting. Rabbit anti-CWLP polyclonal antibody (Ab) was generated by the Biological Services Unit of the Weizmann Institute of Science (Rehovot, Israel) using a polypeptide comprising the last 94 C-terminal amino acid residues of CWLP, which was overexpressed in *E. coli* using pET11a vector (Novagene) and purified by SDS-PAGE. Western blot analyses were performed according to (74). The list of antibodies is depicted in SI, Table S3.

Following transient co-expression of combinations of HA-, GFP-, YFP-, or mCherry-labeled proteins in *N. benthamiana*, the infiltrated leaf samples were subjected to membrane protein extraction as detailed in SI section 8. Membrane proteins were co-immunoprecipitated by polyclonal anti-GFP Ab cross-linked to protein A-agarose beads (Pierce), Chromotek GFP trap, or μMACS™ and MultiMACS™ GFP Isolation Kits (Miltenyi Biotec) according to manufacturers’ instructions. The bound protein complexes were eluted from the beads, resolved by 7.5% gels-SDS-PAGE, and analyzed by immunoblotting as described in SI section 6. To detect CWLP glycosylation, protein extracts from CWLP-GFP seedlings were subjected to immunoprecipitation using immobilized anti-GFP polyclonal Ab. The pulled-down proteins were eluted and size-resolved by SDS-PAGE. After blotting onto nitrocellulose membranes, equivalent blots were stained with Glycoprotein Detection Kit (Sigma-Aldrich) or probed with anti-GFP monoclonal Ab (SI sections 4, 5). For LC-MS/MS mass spectrometry analyses, CWLP-GFP was purified from 3-d-old etiolated CWLP-GFP seedlings, using WT seedling extracts as control and analyzed according to (88) (SI section 7).

### 2.5 Yeast two-hybrid (Y2H) assays

Nuclear two-hybrid interaction assays were performed using the MATCHMAKER GAL4 two-hybrid system (Clontech). The *CWLP* coding region without signal peptide (SP) was cloned in frame downstream of the *GAL4* DNA-binding domain in the bait vector pGBKT7 (Clontech) and introduced into the yeast strain AH109 (Clontech), which was then transformed with a pACT2 carrying cDNA expression library prepared from dark-grown *Arabidopsis* cell suspension (89). Split-ubiquitin Y2H assays were carried out using the DUAL-HUNTER two-hybrid system (DualSystems Biotech). The *CWLP* coding region without SP was cloned in frame between coding sequences of the yeast *invertase* (*SUC2*) signal peptide and *C-Ub* domain in the bait vector pBT3-SUC. Yeast host NMY51 carrying this bait plasmid was transformed with pPR3-N-Arabidopsis cDNA library (DualSystems Biotech Cat. #P02210) as described in SI section 9.

### 2.6 Confocal LSM microscopy

The Olympus IX81/FV500 laser-scanning microscope was used with the following filter sets: for observing GFP, 488 nm excitation and BA505-525 (detector); for Cy3, 543 nm excitation and BA610; for FM4-64, 488 or 515 nm excitation and BA660. We used a PlanApo 60X1.00 WLSM ∞/0.17 objective for live cell imaging and 60 × 1.4 NA PlanApo oil immersion objective for fixed stained samples. When GFP and Cy3 were detected in the same sample, DM (dichroic mirror) 488/543 was used, and sequential acquisition was performed. Transmitted light images were obtained using Nomarski differential interference contrast. Fluorescence intensity was estimated by Image J Software (http://rsb.info.nih.gov/ij). To counterstain the PM and CW, seedlings or detached leaves were incubated respectively in 8 μM FM4-64 (Molecular Probes) and 1.5 μM Propidium Iodide (Sigma-Aldrich) in PBS for 5 or 10 minutes, washed in PBS for a few seconds, and immediately analyzed by confocal microscopy. To immunolocalize proteins, antibodies were introduced into the cells by freeze-shattering. Leaves were immersed in PME buffer and processed according to (90). Probing was performed using either rabbit anti-CWLP or mouse anti-tubulin antibodies, which were then detected by Cy3-goat anti-rabbit IgG or Alexa Fluor 488-goat anti-mouse IgG (Table 3), respectively, as described in SI, Section 2.

### 2.7 Bioinformatics and statistics

Details of applied bioinformatic analyses are described in SI, Section 10. All experiments and analyses were repeated at least three times. Data presented as mean ± SE. P values (*p < 0.05, **P < 0.01, ***P < 0.001) were generated with two-tailed Student t-test paired samples. Pearson’s correlation coefficient was used for co-localization significance when ρ > 0.5.

### 2.8 Accession Numbers

*CWLP* (At3g22120), *DRP1A* (At5g42080), *DRP1C* (At1g14830), *EARLI1* (At4g12480), *FLOT 2* (At5g25260), *GF14* (At1g35160), *PIP2;7/PIP3A* (At4g35100), *PIP2;1/PIP2A* (At3g53420), *PIP1;3/PIP1C* (At1g01620), *PP2A-B’’α* (At5g44090), *PP2A-B’’β* (At5g28850), *PP2A-B’’γ* (At5g28900), *PP2A-B’’δ* (At1g54450), *PRP940* (At3g10940).

## 3 RESULTS

### 3.1 Expression patterns, structural features, and CW-PM localization of CWLP and PRP940

From the four Arabidopsis HyPRPs classified into the C-Type subfamily based on 68 to 80% amino acid identity among their 8CM domains [Fig.1A; (1)], expression of *CWLP* and *PRP940* has been detected in shoot and reproductive organs. By contrast, transcription of *At1g62500*/*PRP14* and *At4g15160*/*PRP17* was observed in young roots and mature root apices according to TAIR (https://www.arabidopsis.org), BAR (https://bar.utoronto.ca), and Genevestigator (https://genevestigator.com) transcript profiling databases. These data were supported by monitoring the activity pattern of the GUS reporter gene fused to 2kb *CWLP* promoter region extending 25 bp downstream of transcription start in homozygous T3 Arabidopsis seedlings (Fig. S1A). pCWLP-GUS reporter activity was detected in cotyledons, leaves, stems, flowers, and siliques, but not in roots (except for root apices, Fig. S1A) in agreement with transcript profiling data indicating *CWLP* expression in seedlings, young leaves, shoots and their meristematic tissues (Fig. S1B, middle panel). *CWLP* is transcribed bi- directionally, producing a reverse complementary long non-coding RNA (*lncRNA At3g22121*), which terminates 132 nucleotides downstream of *CWLP* transcription start site. The double-stranded RNA forming region of 852 nucleotides produces natural antisense small interfering RNAs (NAT-siRNAs) in a phased fashion (91, 92). According to Genevestigator, low levels of *lcRNA* are present in seeds, xylem, and root pericycle tissues (Fig. S1B, right panel), where nat-siRNAs might be expected to modulate *CWLP* expression. In comparison, mRNA levels of *PRP940* are notably higher than those of *CWLP* in areal parts of plants (Fig. S1B, left panel). Our qRT-PCR measurements (Fig. S1C) confirmed that *PRP940* mRNA levels in rosette leaves of seedlings were about 2.5 to 10-fold higher throughout different stages of vegetative development. Remarkably, Genevestigator co-expression analyses indicated that *PRP940* is the top-scoring gene showing positive co-regulation with *CWLP* (Pearson’s correlation coefficient 0.53) as well as that, similarly to *PRP40*, transcription of *CWLP* is negatively modulated by drought, heat, ABA, flagellin 22 and other pathogenic stimuli but stimulated to higher level in response to cold-stress (Fig. S1D). In comparison, (93) detected a slight reduction of *CWLP/lcRNA*-derived nat-siRNA levels in response to drought and cold, in contrast to a minimal increase under salt stress, while transcript profiling databases indicate no significant expression of *lncRNA* under abiotic stress conditions (Fig. S1D).

**Figure 1.**
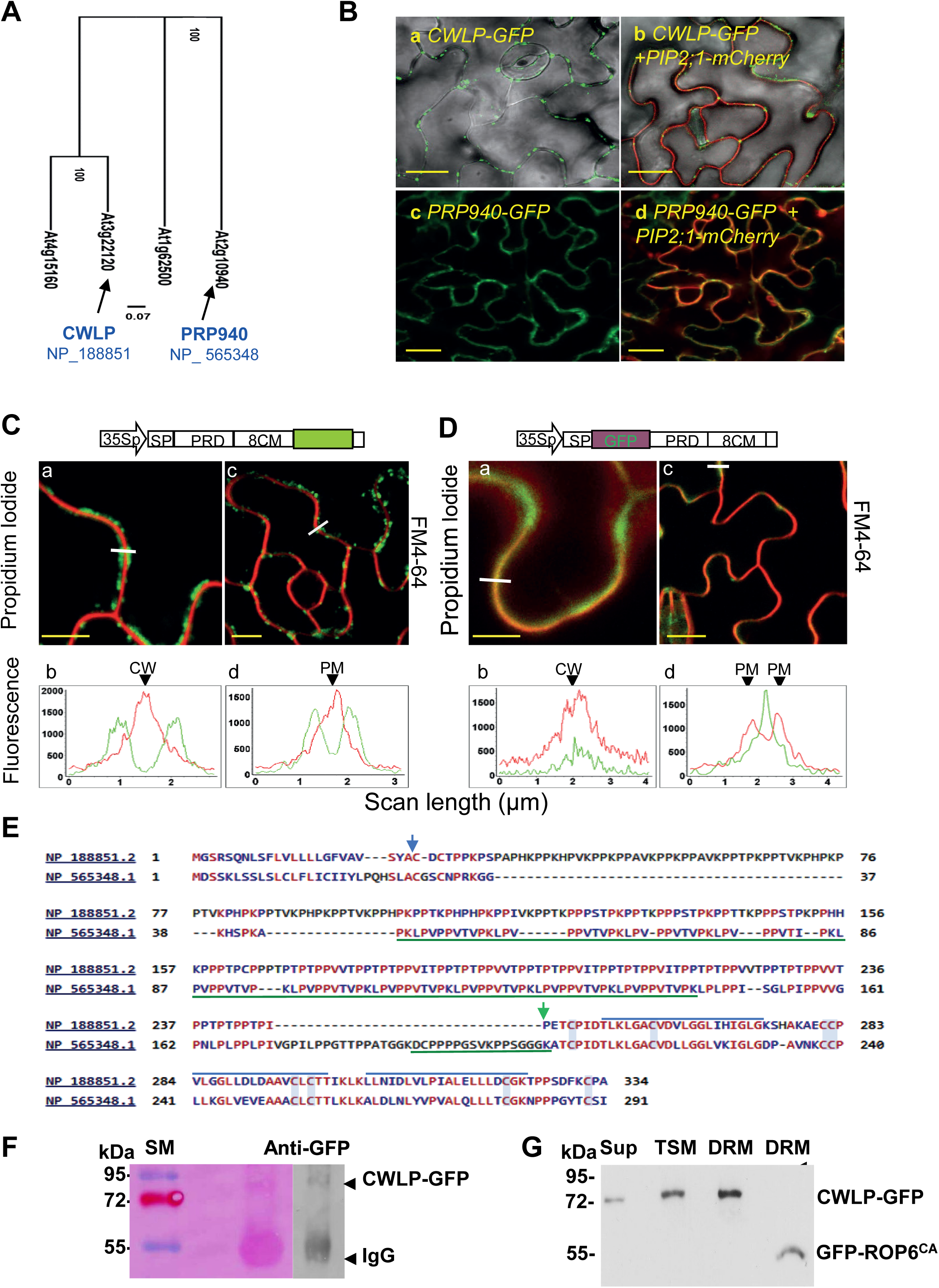
CWLP and PRP940 are closely related. (A) Phylogenetic relationships according to protein sequence homology among the C-type HyPRP proteins in *Arabidopsis*. BLOSUM62 algorithm (Mafft) and RAxML were used for multiple sequence alignment and phylogenetic analysis, respectively. (B) CWLP and PRP940 are localized to the PM, similarly to aquaporin PIP2;1 (At3g53420), which served as a PM marker. Co-localization in leaf epidermal cells of CWLP-GFP (a) and CWLP-GFP with PIP2;1-mCherry (Pearson R value = 0.83) (b), stably expressed in transgenic Arabidopsis plants using the CaMV 35S promoter (35Sp). Bar = 20 μm. PRP940-GFP is similarly localized to the PM (c, d) and co-localized with PIP2;1-mCherry (Pearson R value = 0.87) in *N. benthamiana* epidermal cells during transient expression (d). Bar = 10 μm. (C) and (D) CWLP is localized to the PM with N- and C-termini oriented towards the CW and cytosol, respectively. Leaf epidermal cells of CWLP-GFP (C) and GFP-CWLP (D) expressing plants stained with propidium iodide (PI, red) or FM4-64 (red). Fluorescence peaks of GFP, PI, and FM4-64 (shown in C b, d and D b, d) were obtained by scanning along the white lines indicated in Ca, Cc and Da, Dc. Schemes of expressed gene cassettes are shown above the corresponding images. SP – signal peptide, PRD – proline-rich domain, 8CM – eight-cysteine motif/transmembrane domain. (E) CWLP (NP188851.2) and PRP940 (NP565348.1) sequence alignment indicates common composition of N-terminal signal peptide ending at the blue arrow, PRP domain with reiterated motifs spanning till the green arrow, and 8CM domain with a conserved arrangement of 8 Cys residues marked in blue. Identical amino acid residues are highlighted in red. Blue lines above the CWLP sequence indicate its predicted transmembrane motifs (Fig. S1E), and green lines under the PRP940 sequence show two regions of redundant sequences containing hydroxyproline residues identified by MS/MS peptide analysis performed by (50). (F) CWLP is glycosylated. CWLP-GFP was immunoprecipitated using a polyclonal anti-GFP Ab. The pulled-down CWLP-GFP sample was separated by SDS-PAGE (10% gel) and blotted onto a nitrocellulose membrane. The left half was stained for glycosylation, and the right half was probed with an anti-GFP monoclonal Ab. SM, size markers. (G) CWLP is a membrane protein, mostly present in Triton X-100 soluble (TSM) and detergent-resistant (DRM-nanodomain) membrane fractions extracted from 15-d-old seedlings overexpressing either CWLP-GFP or GFP-ROP6^CA^. A relatively lower amount of CWLP was detected in the supernatant (Sup) following membrane isolation by 100,000 g centrifugation. Western blot analysis of equal-volume samples of each fraction probed with anti-GFP Ab.

Published studies suggested that N-terminal PRDs of BnPRP and CWLP extend towards the CW, whereas their C-termini are oriented towards the PM and cytosol (65, 67). Our bioinformatic studies using structure prediction programs (Fig. S1E to G) indicated that PRD of CWLP forms intrinsically disordered structures. Hydropathy analyses predicted that the C-terminal 8CM region of CWLP might form three transmembrane domains (Fig. S1F). The 8CM domains of BnPRP, CWLP, and PRP940 showed the highest structural similarity to those of Hydrophobic Protein from Soybean [PDB entry 1HYP; (94)] and nsLTPs (59, 95). ConSurf modeling [Fig. S2A and B, (96)] confirmed that, like 1HYP, CWLP 8CM forms a tunnel-like hydrophobic α-helical lipid-binding cavity, which is stabilized by disulfide-bridges between 8CM Cys 1–5, 2–3, 4–7, and 6–8 residues as in 8CMs of snLTP2s. Recent mass spectrometry data suggest potential S-acylation of CWLP at position Cys349 (Cys8 of 8CM) and PRP940 at multiple Cys residues [Cys 1-4, 6 and 8 of 8CM; (97)]. The 8CM domain thus defines a lipid-interacting PM-anchor for CWLP and PRP940.

To visualize the cellular localization of CWLP and PRP940, we generated transgenic Arabidopsis plants expressing C-terminal translational fusions of CWLP with GFP, CFP, HA or mCherry, PRP940-GFP, and GFP-CWLP, harboring GFP between the N-terminal signal peptide and PRD. Confocal microscopy studies using aquaporin PIP2;1 (Plasma Membrane-Intrinsic Protein 2;1) fused to mCherry as PM marker (98, 99) localized both CWLP-GFP and PRP940-GFP to the PM of leaf epidermal cells (Fig. 1B). CWLP-GFP in Arabidopsis, and PRP940 in *N. benthamiana* leaves showed overlapping localization with PIP2;1-mCherry in PM-clusters, named “puncta-like structures” by (100; Fig. 1B). Using ectopically expressed N- and C-terminal GFP fusions, the polar orientation of CWLP termini was examined in abaxial epidermal cells of rosette leaves counter-stained with red CW and PM dyes propidium iodide and FM4-64, respectively. CWLP-GFP was observed in patches at the intracellular surface of CW and PM (Fig. 1Ca and Cc, respectively). The signal of CWLP-GFP patches was detected across the boundaries of adjacent cells (Figs. 1Cb and Cd). This indicated that the C-terminal GFP tag is located in the cytosolic surface of the PM. By contrast, “punctate clusters” of GFP-CWLP overlapped with the PI-stained CWs (Figs. 1Da and Db). Remarkably, when the PM was stained with FM4-64, the fluorescence scanning trajectories across adjacent cells showed red-fluorescent peaks on both sides of the GFP green peak (Fig. 1Dc,d), indicating that the CWLP N-terminus is localized to the vicinity of CW. These observations demonstrated that the CWLP as a linker protein establishes a continuum between CW and cytosol through the PM, as was previously speculated (65, 67, 70).

### 3.2 Verification of glycosylation, nanodomain anchoring, and cell wall linker function of CWLP

Both CWLP and PRP940 were identified in CW protein fractions by mass spectrometry, revealing the presence of HyP residues in tryptic peptides of PRP940, which show considerable conservation in the PRD repeats of CWLP [underlined in green, Fig. 1E; (50)]. To confirm predicted O-glycosylation of HyP residues, protein samples from total cell-extracts of CWLP-GFP-expressing plants were immunoprecipitated with polyclonal anti-GFP Ab and subjected to western blotting using glycosylated IgG as positive control (Fig. 1F, 55 kDa), followed by visualization with Glycoprotein-Detection Kit and monoclonal anti-GFP antibody (Fig. 1F). CWLP-GFP (64.7 kDa) was detected as weakly glycosylated protein migrating with an apparent molecular mass of 90 kDa in SDS-PAGE (Fig. 1F). Glycosylation of CWLP and likely PRP940 HyP residues could thus mediate N-terminal interaction of these proteins with polysaccharide components of the CW as suggested (3).

By using single-particle tracking, Li, *et al* (98) demonstrated that aquaporin PIP2;1 can move in and out of PM nano/microdomains (99). Co-localization of CWLP and PRP940 with PIP2;1 (Fig. 1B) in PM clusters thus suggested that CWLP and PRP940 are localized in PIP2;1-marked PM-NDs, which are composed of “detergent-resistant membranes” (DRMs) that cannot be entirely solubilized by the nonionic detergent Triton X-100 at low temperature (101–103). To confirm that CWLP is a DRM-associated protein, total cell extract from CWLP-GFP-expressing 15-d-old seedlings was fractionated to cleared (100,000 g) supernatant (Sup), Triton X-100 solubilized (TSM), and DRM protein fractions and analyzed by western blotting using anti-GFP Ab (Fig. 1G). As a control, similar protein fractions were prepared from GFP-ROP6^CA^ (DRM-localized mutant of Rho-related-GTPase) expressing plants grown in parallel under identical conditions (87). CWLP-GFP was detected in all three cell fractions, including DRM (Fig. 1G). This observation indicated that part of CWLP is present in nanodomain-derived DRMs, but also detectable in the TSM and Sup fractions, likely due to the dynamic equilibrium between PM-trafficking, clustering, and recycling of these nanodomain proteins (100, 101, 103–105).

Exposure of plant cells to a hypertonic environment results in plasmolysis, including protoplasm shrinkage and detachment of PM from the CW, sometimes retaining only few adhesion sites known as “Hechtian strands” (106). We employed a plasmolysis assay to verify CWLP functions as a genuine CW-PM linker. Detached leaves of 14-d-old Arabidopsis seedlings overexpressing CWLP-GFP were immersed for 20 min in 30% glycerol (Fig. 2Ad-f) or water as control (Fig. 2Aa-c), and protoplasm shrinkage of epidermal cells was compared to similarly treated leaves of WT (Figs. 2Aa and Ad) and GFP-overexpressing (Fig. 2Ab and Ae) cells. While WT and GFP-OE control cells responded with pronounced protoplasm shrinkage to glycerol treatment (Fig. 2Ad and 2Ae, respectively), no separation of PM and CW was detectable in CWLP-GFP expressing cells (Fig. 2Af). The separation of CW and PM during hyperosmotic stress is accompanied by the destruction of the microtubule network (107). In plasmolysis assays combined with immunolocalization of tubulin, we observed that protoplast shrinkage induced by 30% glycerol destroyed the microtubule network in WT control (Fig. 2Ba,b and Bc). No plasmolysis and disassembly of microtubule cytoskeleton were detected in similarly treated CWLP-GFP (Fig. 2Bd-g) and CWLP-HA (Fig. 2Bh-k) overexpressing leaf epidermal cells. Similar plasmolysis-tolerance was observed in leaf cells of CWLP-GFP-OE seedlings treated with 0.6 M sorbitol for 30 minutes (Fig. 3A, Cb).

**Figure 2.**
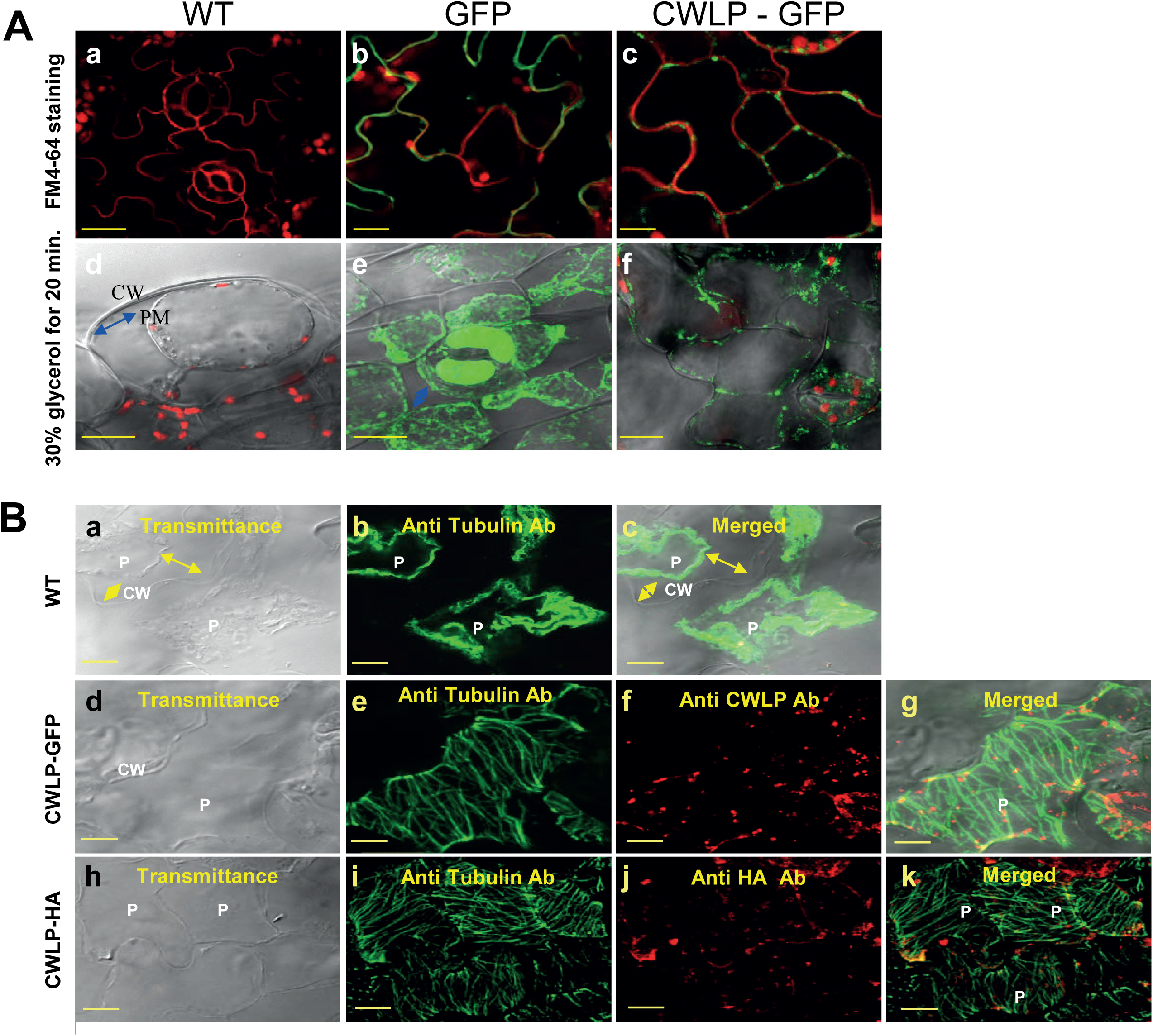
*Arabidopsis* leaves overexpressing CWLP-GFP or CWLP-HA show enhanced tolerance to hyperosmotic stress. (A) Comparison of the response to 30% glycerol of leaf epidermal cells of 14-d-old plants: (**a to c)**: Epidermal cells of excised leaves immersed in water and stained with FM4-64 (Red) of Wild Type (a), GFP overexpressing (b), and CWLP-GFP overexpressing (c) plants. GFP is present in peripheral cytoplasm (b), CWLP-GFP displays dotted PM localization (c). (**d to f**): Immersion of excised leaves in 30% glycerol for 20 min. Plasmolysis occurred in epidermal cells of WT (d) and GFP overexpressing (e) leaves, where GFP is localized in the cytoplasm. CWLP-GFP leaves (f) did not show any plasmolysis or change in the dotted PM localization. The blue arrow in (d) shows protoplasm shrinkage. CW – cell wall, PM – plasma membrane. Scale bars are 20 μm in a, d, e, f, and 10 μm in b and c. (B) Response of microtubule cytoskeleton to plasmolysis. Overexpression of CWLP-GFP or CWLP-HA prevents destruction of the microtubule network. Detached leaves of 14-d-old plants were exposed to 30% glycerol for 20 minutes, fixed, and subjected to freeze-shuttering, allowing cellular penetration of anti-CWLP, anti-tubulin, and anti-HA primary antibodies as indicated, and then secondary fluorescent antibodies conjugated to Cy3 (red) for CWLP or HA, and to Alexa fluor 488 (green) for tubulin detection. Z-stack images are presented. (**a to c**): Plasmolysis in cells of WT plants. Disrupted tubulin-cytoskeleton structure of plasmolyzed cells is displayed by comparison of transmittance (a), tubulin fluorescence (b), and merged (c) images. Arrows indicate the gap between the contracted protoplasm (P) and CW. (**d to g**): Intact cytoskeleton of 30% glycerol-treated, non-plasmolyzed cells of CWLP-GFP leaves, visualized by transmittance (d), tubulin (e), and CWLP (f) fluorescence, and merged (g) images. (**h to k**): No plasmolysis and maintaining of the intact cytoskeleton in 30%-glycerol-treated CWLP-HA cells were detectable by transmittance (h), tubulin fluorescence (i), CWLP-HA fluorescence (j), and combined image (k) analyses. CW- Cell wall, P- Protoplasm, Bars = 10 μm.

**Figure 3.**
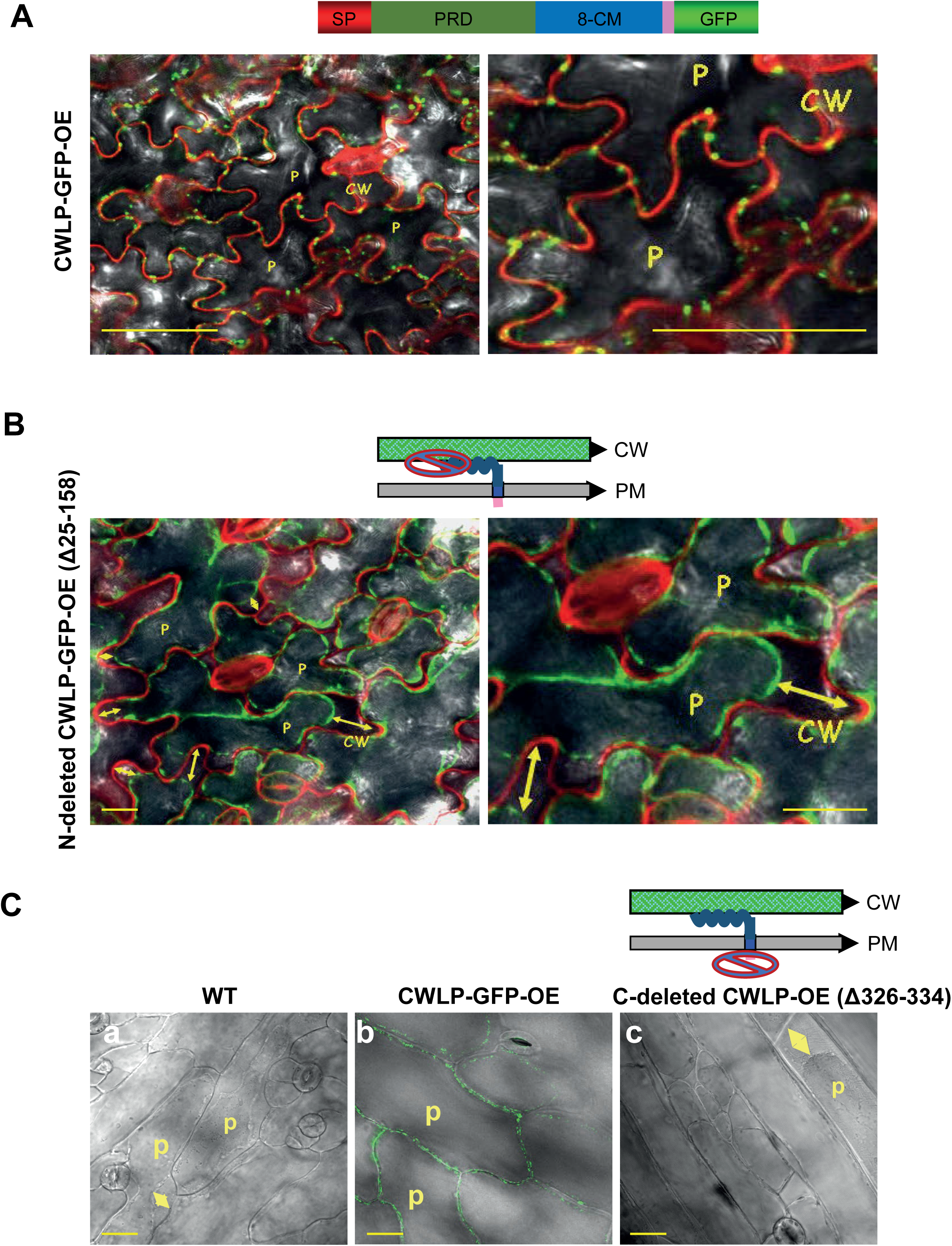
C- and N-termini of CWLP are essential for enhanced plasmolysis tolerance. Plasmolysis assays were performed by exposing detached leaves of 14-d-old plants to 0.6M sorbitol for 30 min. (A) No plasmolysis in epidermal cells of plants overexpressing the full-length CWLP-GFP control construct. Bar = 50 μm. The magnified image is displayed in the right panel. Bar = 10 μm. The upper scheme displays domains of the full-length CWLP-GFP. (B) Plasmolysis in epidermal cells of plants overexpressing CWLP-GFP (Δ25-158) carrying a deletion in PRD. CW is stained in red with PI. Bar = 20 μm. Magnified image is displayed in the right panel. Bar = 10 μm. (C) Plasmolysis in abaxial epidermal cells, which are close to the leaf midrib in plants overexpressing CWLP (Δ326-334) lacking 8 C-terminal amino acid residues (c), compared with plasmolysed WT (a) and non-plasmolysed CWLP-GFP (b) cells. Bars= 20 μm. Upper schemes display the deletions in B and C. Yellow arrows indicate PM detachment from CW, P-protoplasm.**c d**

According to structural studies described above, the N-terminus of CWLP is likely linked to the CW by its glycosylated PRD, whereas its C-terminal 8CM transmembrane domain serves as a PM anchor. To examine how the removal of CWLP termini affects the CW-PM linker function, we deleted 133 amino acid residues downstream to the SP in the PRD domain of CWLP-GFP (Δ 25-158), or the last eight amino acids of the C-terminal 8CM domain (Δ326-334) in CWLP. Excised leaves of homozygous T3 transgenic plants overexpressing these constructs were immersed in 0.6 M sorbitol for 30 min or in water as control. Compared with CWLP-GFP cells that showed no PM detachment from the CW (Fig. 3A), cells overexpressing the N- or C-terminally truncated versions of CWLP-GFP (Fig. 3B) or CWLP (Fig. 3Cc) respectively, underwent plasmolysis similarly to WT and GFP-expressing plants (Fig. 2A). Thus, both CWLP termini are essential for maintaining CW-PM continuum and tolerance to plasmolysis,

### 3.3 Role of actin-myosin network in PM-ND localization of CWLP and its association with dynamin-related proteins

Cytoskeletal networks of microtubules (MTs) and actin filaments (AFs) have distinct and interplaying roles in peripheral export (exocytosis) and recycling (endocytosis) of various cell wall and PM components (29, 108). To test the potential association of CWLP with MTs or AFs, we transiently co-expressed CWLP-GFP with the microtubule-binding domain of CASEIN KINASE1-LIKE-6 fused to mCherry [CKL6-CTD-mCherry; (109)] and a DsRed-tagged version of AF-binding FIMBRIN-1 protein AtABD2-DsRed (110) in *N. benthamiana* leaves, respectively. Using quantitative scanning for overlapping fluorescent signals, we did not observe co-localization of CWLP-GFP clusters with MTs and AFs (Fig. 4A and B). Therefore, we next examined whether oryzalin and latrunculin B (LatB), which disrupt the MT and AF networks, respectively, would alter clustered localization of CWLP-GFP in PM-NDs. Leaves of 14-d-old Arabidopsis plants expressing either α-TUBULIN6-GFP or CWLP-GFP were simultaneously incubated with oryzalin and then stained with the PM dye FM4-64 (Fig. 4C). The α-TUBULIN6-GFP-labelled MT-network (green filaments in Fig. 4Ca) was destroyed by oryzalin treatment as expected (Fig. 4Cb), but oryzalin did not change clustered PM-localization of CWLP-GFP (overlap of punctate green (Fig. 4Cc) and red PM (Fig. 4Cd) patterns). Similarly, oryzalin treatment did not alter the epidermal cell localization pattern of PRP940-GFP (Fig. S3). On the other hand, LatB treatment of Arabidopsis leaves expressing either the actin-binding protein mTalin-GFP (111) or CWLP-GFP, followed by staining with FM4-64, showed that LatB completely diminished mTalin-GFP-labeling of dissociated AF network (Fig. 4Da and Db) and enhanced internalization of CWLP-GFP in endocytic vesicles (Fig. 4Dc and Dd, white arrow), corroborating that functional AF network is essential for regular CWLP assembly (9). While the shifting of CWLP-GFP to endosomes caused by LatB treatment also confirmed that LatB does not interfere with the initiation of PM-ND proteins internalization, and the formed endosomes are visually prominent because LatB prevents their further cellular trafficking and dynamic recycling (40).

**Figure 4.**
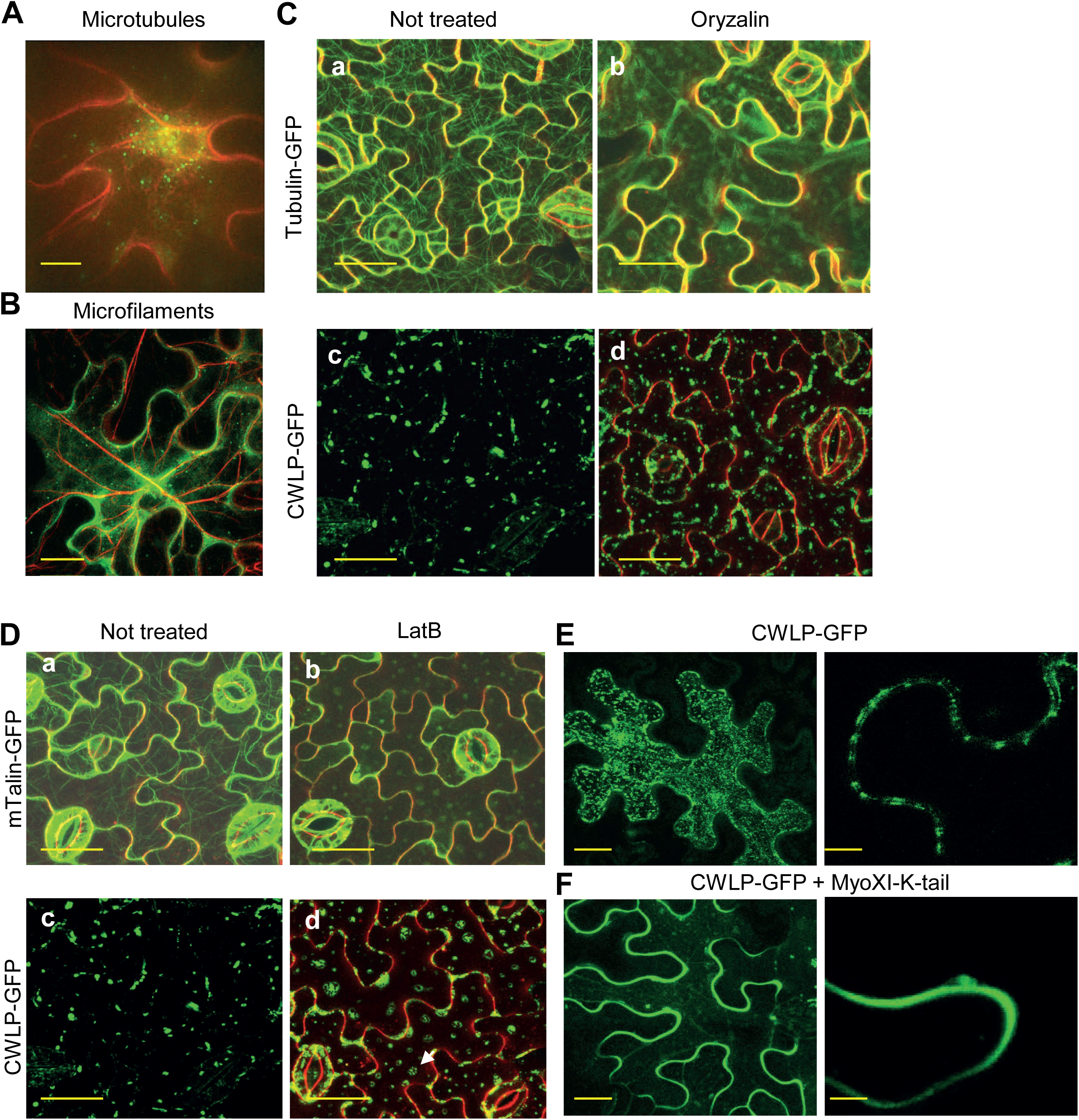
Correlation between PM-localization of CWLP-GFP and intactness of cytoskeleton in leaves. (A and B) CWLP-GFP does not show co-localization with microtubules and actin microfilaments. Spinning disc confocal microscope images of *N. benthamiana* epidermal cells transiently co-expressing CWLP-GFP with CLK6-CTD-mCherry that binds tubulin (A), or with Fimbrin1-ABD2-DsRed, which interacts with microfilaments (B). Bar = 50 μm in (A) and 20 μm in (B). (C) Intact microtubules are not required for PM localization of CWLP-GFP. (a) Microtubule network displayed in epidermal cells of transgenic *Arabidopsis* expressing tubulin α6-GFP. (b) Destruction of microtubule network, shown in (a), by oryzalin treatment. (c) Distribution pattern of CWLP-GFP in the PM of *Arabidopsis* leaf epidermal cells. (d) Oryzalin treatment does not alter PM localization of CWLP-GFP in leaf epidermal cells. Red – PM staining with FM4-64. Bar = 20 μm. (D) Intact AFs are essential for internalization CWLP-GFP. (a) Microfilament network displayed in leaf epidermal cells of transgenic *Arabidopsis* expressing mouse mTalin-GFP. (b) LatB (25μM for 2 h) eliminates the microfilament network shown in (a). (**c and d**) PM-localization of CWLP-GFP shown in (c) is interrupted by LatB leading to CWLP internalization as endocytic vesicles (white arrow). Red – PM staining with FM4-64. Bar = 20 μm. (E and F) Blocking the actomyosin mediated cytoskeleton trafficking by overexpressing dominant negative MyoXIK tail-domain interferes with PM-nanodomain clustering of CWLP-GFP in *N. benthamiana* epidermal cells. (E) Untreated control showing normal clustering of CWLP-GFP. (F) Lack of CWLP-GFP clustering due to co-expression of CWLP-GFP with the mutated MyoXIK tail-domain. Bar = 20 μm in left panels and 5 μm in right panels.

Next, we examined how LatB-treatment affects co-localization of CWLP-GFP and aquaporin PIP2;1-mCherry in the PM. In addition to detecting parallel internalization of aquaporin, LatB-treatment resulted in significant enlargement and extension of overlapping PM-domains of CWLP-GFP and PIP2;1-mCherry (Fig. S4, orange PM sections), while only a fraction of PIP2;1-mCherry showed co-localization with CWLP-GFP in the endocytic vesicles.

Members of the *Arabidopsis* myosin XI family play essential roles in organellar and vesicular trafficking. Overexpression of myosin XI-K-tail domain lacking two functionally important arginine residues was reported to block actomyosin-mediated cellular and PM trafficking (112). Transient co-expression of mutant myosin XI-K tail domain with CWLP-GFP in *N. benthamiana* leaves abolished PM-ND clustering of CWLP-GFP (Fig. 4E), resulting in even distribution of CWLP-GFP along the PM (Fig. 4F), confirming that a functional actomyosin cytoskeleton is essential for proper targeting and assembly of CWLP clusters in the PM-ND.

Notably, nanodomain localization of CWLP-GFP was not interrupted in leaf protoplasts obtained by CW enzymatic digestion (Fig. S5). This suggested that despite potential CW anchoring of glycosylated CWLP-PRD (Fig. 1Da and Db) and its requirement for conferring plasmolysis tolerance (Fig. 3B), CW digestion does not affect CWLP assembly in PM-NDs. However, the involvement of the C-terminus in CWLP proper functionality suggested that the CWLP and its cargo partners might be recognized early on (i.e., already upon its translation in the ER) by components of the AF network, which assist its subsequent trafficking to PM-NDs.

In order to identify CWLP-associated proteins, cytosolic and membrane protein fractions were prepared from etiolated WT (control) and CWLP-GFP expressing seedlings. Subsequently, CWLP-GFP was affinity purified from the cytosolic fraction by immobilized anti-GFP antibody. Following separation of eluted proteins by SDS-PAGE (Fig. 5A), the protein bands were excised, and their tryptic peptides were analysed by LC-MS/MS. Four proteins were identified: BIP1 (band 1, 73.6 kDa, At5g28540) and BAG7, a Bcl-2-associated athanogene protein (band 4, 51.6 kDa, At5g62390), involved in maintaining protein unfolding in the ER during cold or heat exposure (113, 114), as well as DYNAMIN-RELATED PROTEINS DRP1C (AT1G14830) and DRP1A (AT5G42080) (bands 2 and 3; Fig. 5B) at a peptide coverage of 78% and 74%, respectively (SI, Table S1). Western blot analysis of GFP-trap purified CWLP-GFP complexes using anti-DRP1A or anti-DRP1C antibodies (115, 116) confirmed association of CWLP with both DRP1A and DRP1C *in vivo* (Fig. 5C, lanes 6 and 8), which was absent in the WT control (Fig. 5C, lanes 5 and 7). DRP1A and DRP1C are involved in the scission and release of endocytic PM vesicles (117–119), directly bind to AFs at the PM, and play a pivotal role in post-Golgi vesicular trafficking in both CME and CIE pathways (33, 120, 121). Consequently, our mass spectrometry data indicated that CWLP likely establishes a link to the AF network by interacting with DRPs.

**Figure 5.**
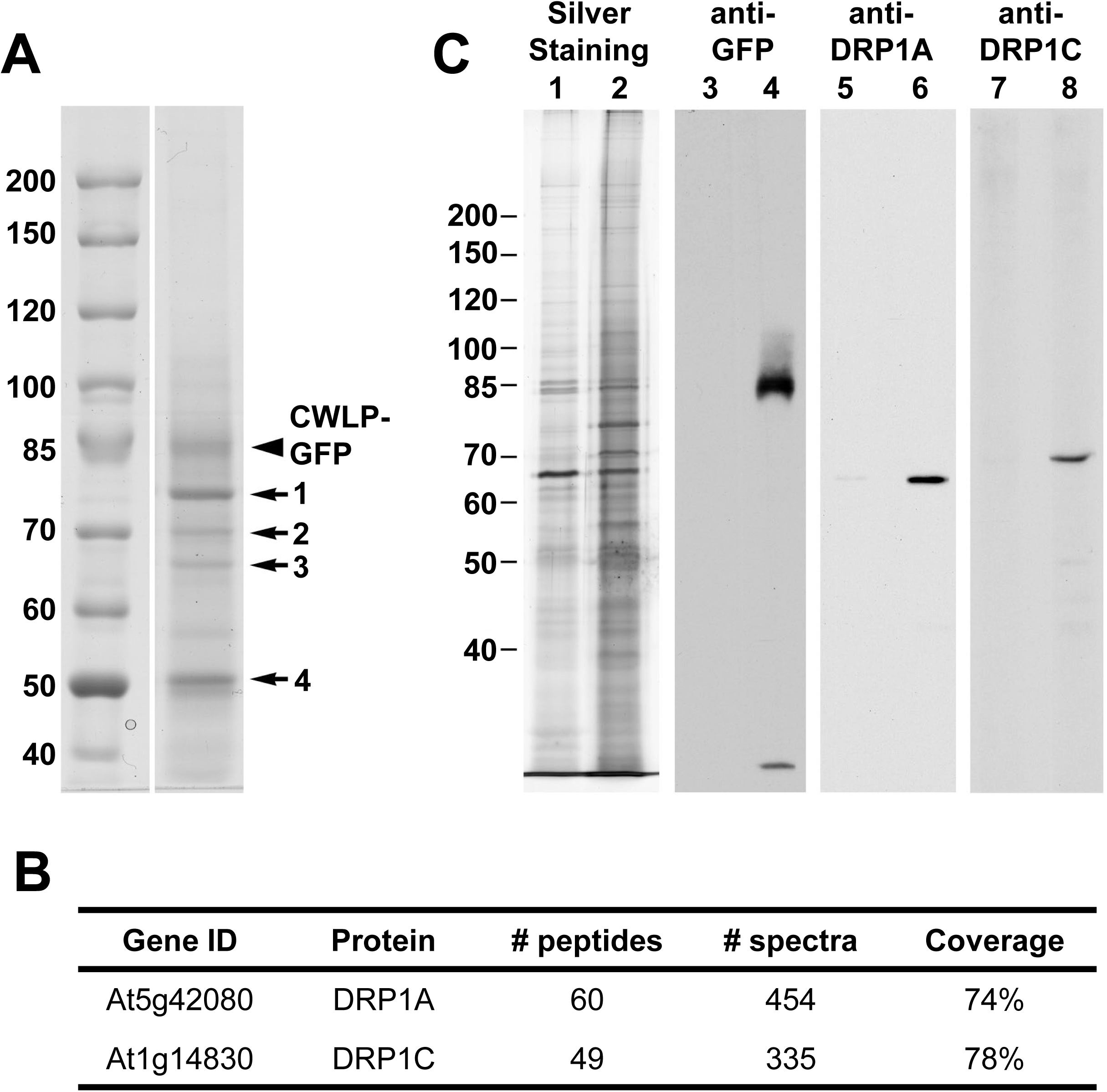
LC-MS/MS analysis of CWLP-GFP associated proteins in the cytoplasmic fraction. (A) Size fractionation of CWLP-GFP-associated proteins eluted from GFP-Trap by SDS-PAGE. In addition to CWLP-GFP, four protein bands (1-4) were excised and subjected to identification of their tryptic peptides by LC-MS/MS. Molecular weight markers in kDa are shown on the left. (B) Identification of DRP1C (band 2) and DRP1A (band 3) by peptide analysis (SI Table 1 online). (C) Silver staining and immunoblot analysis of GFP-Trap purified protein fractions from wild-type (control) and CWLP-GFP-overexpressing plants using anti-GFP, DRP1A, or 1C antibodies. Lanes 1, 3, 5, and 7, control-IP; lanes 2, 4, 6, and 8, CWLP-GFP-IP.

### 3.4 Aquaporin binding and recruitment of PP2A phosphatase B’’-subunits by CWLP and PRP940 suggest a role for PP2A in the dephosphorylation-triggered closure of aquaporin water channels

DRP1A, DRP1C, and several other dynamin-related proteins have also been identified by mass spectrometry in purified cytoplasmic fractions of aquaporins PIP1;2 and PIP2;1 (Table 3 in 122). To examine whether CWLP directly interacts with aquaporins, we used CWLP without its signal peptide as bait in yeast split-ubiquitin two-hybrid (Y2H) screens of Arabidopsis cDNA library (123). CWLP was found to interact with aquaporins PIP2;7/PIP3A (At4g35100), PIP2;1/PIP2A (At3g53420), and PIP1;3/PIP1C (At1g01620), from which PIP2;7 conferred the highest level of lacZ-reporter activation in yeast (Fig. 6A). Remarkable structural conservation of PIP2;7 illustrated by ConSurf analyses (96) predicted that CWLP might also interact with other aquaporins.

**Figure 6.**
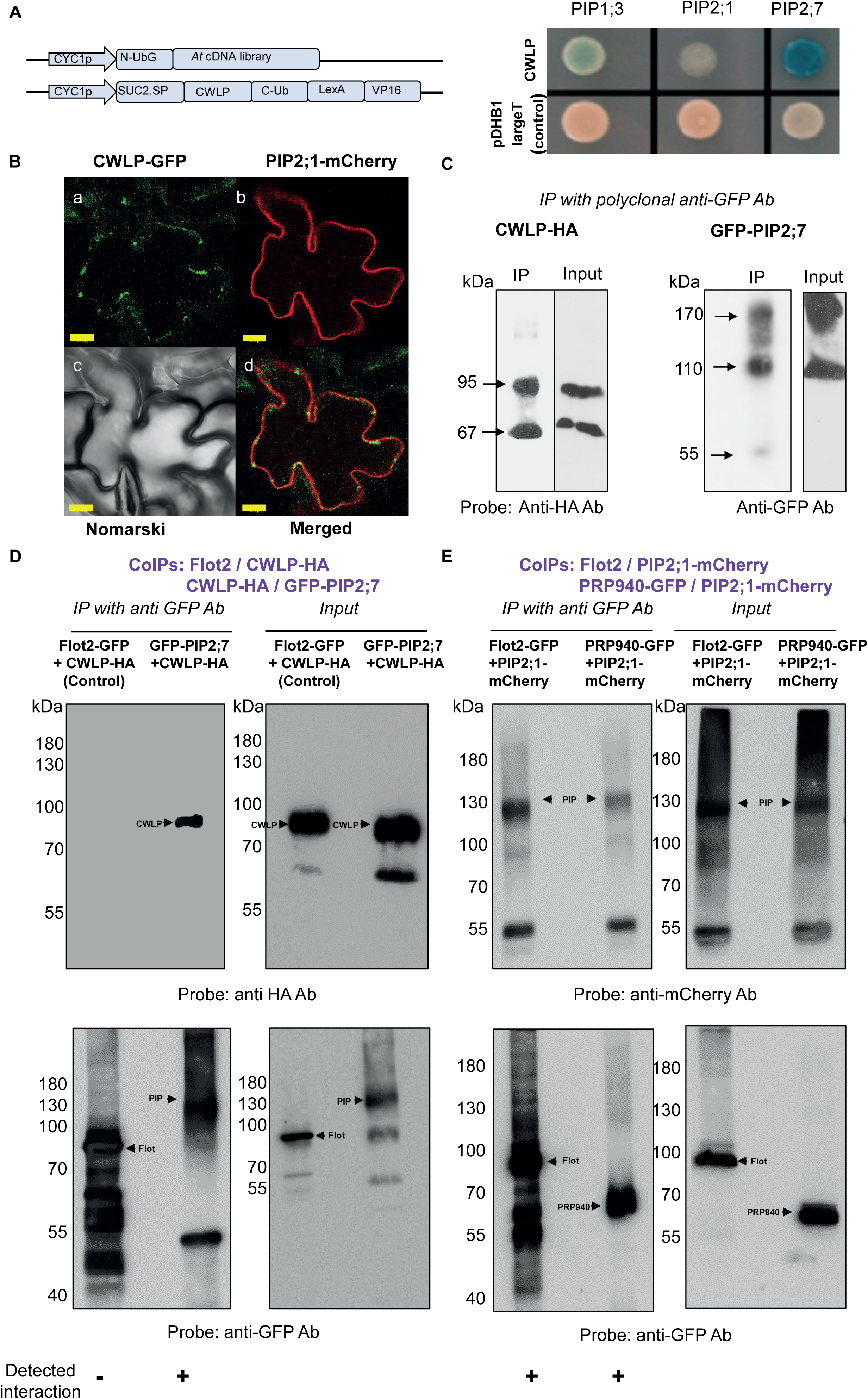
CWLP and PRP940 interact with PIPs, whereas PIP2;1 interacts with the nanodomain marker Flot2. (A) CWLP interacts with aquaporins in split-ubiquitin Y2H assays: the yeast NMY51 strain was co-transformed with pPR3-N prey vector harboring *Arabidopsis* cDNA (upper construct). pBT3-SUC, harboring the *CWLP* coding region lacking the signal-peptide sequence (*CWLP*), downstream of the yeast *SUC2* signal peptide, served as bait (lower construct). Interactions with three aquaporins were identified by monitoring lacZ activity in colonies growing on a selective medium with X-Gal (right panel). (B) Representative image of transient co-expression of CWLP-GFP and PIP2;1-mCherry in leaf epidermis cells of *N. benthamiana* showing a similar pattern of CWLP clustering and co-localization with PIP2;1 as was detected by stable co-expression of both proteins in transgenic *Arabidopsis* plants (Fig. 1Ba, b). Bar = 10 μm. (C) Co-IP of GFP-PIP2;7 with CWLP-HA, indicating mutual interaction in the PM. Membrane proteins from *N. benthamiana* leaves, transiently co-expressing CWLP-HA and GFP-PIP2;7, were isolated and immunoprecipitated (IP) using anti-GFP polyclonal Ab cross-linked to protein A-agarose beads. Precipitation with protein A-agarose beads lacking Ab was used as a negative control. Western blots probed with anti-HA (left panel) or anti-GFP (right panel) monoclonal Abs identified co-IP of CWLP-HA and GFP-PIP2;7 (IP lanes). Like many membrane proteins (171), CWLP-HA (calculated MW 37.8 kDa) showed slower SDS-PAGE migration and was resolved into two distinct forms with apparent MW of about 65 and 95 kDA suggesting its potential glycosylation and lipid binding by S-acylation. In comparison, GFP-PIP2;7 (56.6 kDa) partly retained the typical tetrameric form of aquaporins (153,172,173) but was also detected as a dimer and low-level monomer. (D) *In planta* interaction of CWLP-HA with GFP-PIP2;7 and no interaction with Flot12-GFP indicating co-IP selectivity. Membrane proteins from *N. benthamiana* leaves, transiently co-expressing the protein-pairs Flot2-GFP with CWLP-HA and GFP-PIP2;7 with CWLP-HA, were isolated and immunoprecipitated using monoclonal anti-GFP trap (Chromotek). Western blots probed with anti-HA (upper panel) or anti-GFP (lower panel) monoclonal Abs revealed co-IP of CWLP-HA with GFP-PIP2;7, but not with Flot2-GFP. (E) *In planta* interaction of PRP940-GFP or Flot2-GFP with PIP2;1-mCherry identified differential interaction in PM-ND. Co-IP of transiently expressed membrane proteins extracted from *N. benthamiana* leaves using anti-GFP trap. Probing western blots with anti-mCherry (upper panel) or anti-GFP-monoclonal Abs identified independent interactions between PRP940 with PIP2;1 and PIP2;1 with Flot2. While revealing only partial dissociation of tetrameric PIP2;1-mCherry, the control anti-GFP western blot indicated slightly shifted migration of FLOT2-GFP (78.3 kDa) and PRP940-GFP (56.6 kDa) membrane proteins in SDS-PAGE.

Given that CWLP-GFP and PIP2;1-mCherry showed similar patterns of co-localization in *Arabidopsis* and *N. benthamiana* leaves (Fig. 1Bb; Fig. 6B), we used the latter system to confirm the association of CWLP with aquaporins. Membranes containing TSM and DRM fractions were isolated from *N. benthamiana* leaves co-expressing CWLP-HA and GFP-PIP2;7 (SI section 8), and affinity-purified on anti-GFP polyclonal Ab matrix. Immunoblots of eluted proteins probed with anti-HA and anti-GFP Abs confirmed co-immunoprecipitation of CWLP-HA with GFP-PIP2;7 (Fig. 6C).

The dimer/oligomer-forming flotillins FLOT1 and FLOT2 were identified by mass spectrometry as abundant marker proteins in purified DRMs [e.g., (21, 22, 124–126)] and demonstrated to co-localize and interact with PIP2;1, PIP2;7 and other aquaporins (43, 98). Therefore, we used co-expression of FLOT2-GFP and PIP2;1-mCherry as a control to test whether PRP940-GFP forms a complex with PIP2;1-mCherry similarly to CWLP when co-expressed in *N. benthamiana* leaves. Upon performing affinity purification on GFP-Trap, immunoblots probed with anti-mCherry Ab revealed co-immunoprecipitation of FLOT2-GFP or PRP940-GFP with PIP2;1-mCherry (Fig. 6E, upper panel), while probing with anti-GFP Ab confirmed the presence of both GFP-tagged bait proteins in the purified complexes (Fig. 6E, lower panel).

Next, we examined whether CWLP occurs in a shared complex with FLOT2 and aquaporin. TSM+DRM protein fractions were prepared from *N. benthamiana* leaves transiently co-expressing either FLOT2-GFP with CWLP-HA or GFP-PIP2;7 with CWLP-HA (Fig. 6D) and purified on GFP-Trap. While anti-HA western blotting detected co-immunoprecipitation of CWLP-HA with GFP-PIP2;7, we did not observe CWLP-HA in association with affinity purified FLOT2-GFP. Similarly, PRP940 did not interact with FLOT2-GFP (Fig. 6E). These results indicated that CWLP and PRP940 form aquaporin complexes independently of FLOT2 in distinct nanodomains and thus revealed a novel class of PM-ND complexes (25, 103, 127).

In complementary yeast 2H screens, CWLP without its signal peptide was fused to the *GAL4BD* and used as bait to find interacting partners encoded by *GAL4AD* fused to clones of Arabidopsis-cell-suspension cDNA library. These screens identified four regulatory B’’ subunits of protein phosphatase-2A [PP2A-B’’α (At5g44090), PP2A-B’’β (At5g28850), PP2A-B’’γ (At5g28900), and PP2A-B’’δ (At1g54450)] as CWLP interactors (Fig. 7A). Heterotrimeric AtPP2A holoenzymes are assembled in various combinations of 5 catalytic (C), three scaffolding (A) and 17 regulatory (B, B’ and B’’) subunits (128). The six AtPP2A-B’’subunits carry EF-hand motifs identified first as Ca^2+^-binding domains in calmodulins (129). Whereas B’’β and B’’γ are almost identical, they share lower sequence homology with other members of the PP2A-B’’subunit family (130). To verify *in planta* PP2A-B’’ recruitment by CWLP, PP2A-B’’γ with a C-terminal *YFP* tag (PP2A-B’’γ-YFP) was co-expressed with CWLP-HA in *N. benthamiana* leaves (Fig. 8C), and membrane proteins were purified by binding to protein A-agarose-bound polyclonal anti-GFP Ab (i.e., recognizes YFP). Immunoblots of eluted proteins probed with anti-HA or anti-GFP Abs indicated that PP2A-B’’γ-YFP (90 kDa; which suffered partial proteolysis during isolation; Fig. 7B, right panel) was associated with the faster migrating (i.e., probably less glycosylated, Fig. 6C) form of CWLP-HA (65kDa; Fig. 7B, left panel). To monitor their interaction in the PM by confocal microscopy, PP2A-B’’γ-YFP was co-expressed with CWLP-CFP in *N. benthamiana* leaves. Overlapping of yellow and cyan fluorescent signals documented co-localization of PP2A-B’’γ-YFP and CWLP-CFP in patchy membrane clusters of different sizes, including some larger ball-shaped ones (Fig. 7C and D, respectively). Intriguingly, when expressed without CWLP, PP2A-B’’γ-YFP was observed in yet undetermined dotted granular structures in the cytoplasm which did not show co-localization with either Golgi or endosomal fluorescent markers (Fig. S6). Thus, it indicated that overexpression of CWLP-CFP enhanced the recruitment of PP2A-B’’γ-YFP to the PM.

**Figure 7.**
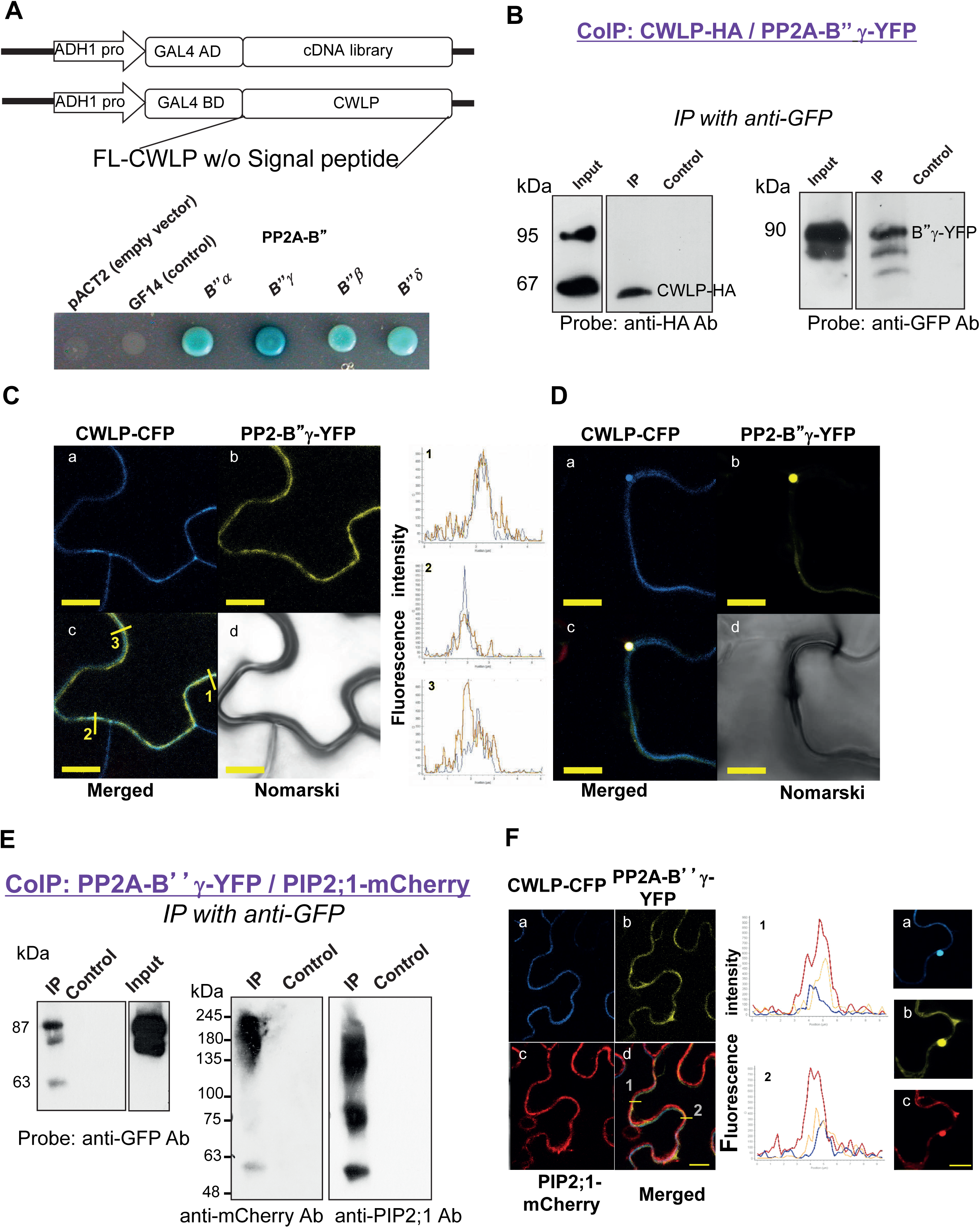
B’’ regulatory subunits of PP2A interact with CWLP and PIPs. (A) Schematic presentation of GAL4-based Y2H nuclear system using *CWLP* cDNA without signal-peptide sequence cloned in pGBKT7 as bait. A library of *Arabidopsis* cDNAs cloned in pACT2 was used as prey. The lower panel shows the interaction between CWLP and four PP2A-B’’ subunits in yeast grown on X-Gal containing medium. GF14 protein served as a negative control. (B) Interaction of CWLP-HA with PP2A-B’’γ-YFP *in planta* demonstrated by Co-IP. Membrane proteins were isolated from *N. benthamiana* leaves transiently co-expressing CWLP-HA and PP2A-B’’γ-YFP. Proteins associated with PP2A-B’’γ-YFP were immunoprecipitated (IP) using protein A agarose cross-linked to polyclonal rabbit anti-GFP Ab and analyzed by western blotting. CWLP-HA and PP2A-B’’γ –YFP were identified by probing with anti-HA (left panel) and anti-GFP (right panel) monoclonal Abs. Input: total membrane fraction before immunoprecipitation. Control: parallel IP using protein A-agarose without cross-linked antibody. (C) Co-localization of CWLP-CFP and PP2-B’’γ-YFP co-expressed in *N. benthamiana* leaves. Single-plane confocal images of CWLP-CFP (a), PP2A-B’’γ-YFP (b), merged (c), and Nomarski (d) images. Co-localization was confirmed by fluorescence measurement (right panel) of the numbered scanned sections indicated by yellow bars in (c) (Pearson R values are 0.54, 0.71, 0.54 for lines 1 to 3, respectively). Blue and yellow lines demonstrate the fluorescence of CWLP-CFP and PP2A-B’’γ-YFP, respectively. Bar = 10 μm. (D) Clusters containing CWLP-CFP and PP2A-B’’γ-YFP are often observed protruding from PM, appearing as ball-shaped membrane bulges. Panels (a to d) are identical to those in (C). Bar = 10 μm. (E) Co-IP of PP2A-B”γ-YFP and PIP2;1-mCherry, indicating interaction *in planta*. Membrane proteins isolated from leaves co-expressing PP2A-B”γ-YFP and PIP2;1-mCherry, were immunoprecipitated using GFP-Trap. Western immunoblotting of the IP fraction identified PP2A-B”γ-YFP by anti-GFP monoclonal antibody (left panel) and PIP2;1-mCherry by both monoclonal anti-mCherry and polyclonal anti-PIP2;7 antibodies (right panel). The anti-PIP2;7 polyclonal antibody better detected partially dissociated forms of PIP2;7 homologs. Input: total membrane proteins prior to the IP. Control: parallel IP using protein A-agarose without cross-linked antibody. (F) Confocal single-plane images of *N. benthamiana* leaf cells co-expressing CWLP-CFP (a), PP2A-B”γ-YFP (b) and PIP2;1-mCherry (c). Merged image of (a), (b) and (c) is shown in (d). Fluorescence scans along the numbered lines in (d) are shown in the middle panel. Blue - CWLP-CFP, yellow - PP2A-B”γ-YFP, and red - PIP2;1-mCherry. Clusters containing CWLP-CFP, PP2-B’’γ-YFP, and PIP2;1-mCherry, protruding from the PM are shown in the right panel (a to c are identical in both panels), indicating co-localization of the three interacting proteins. Bars = 10 μm.

**Figure 8.**
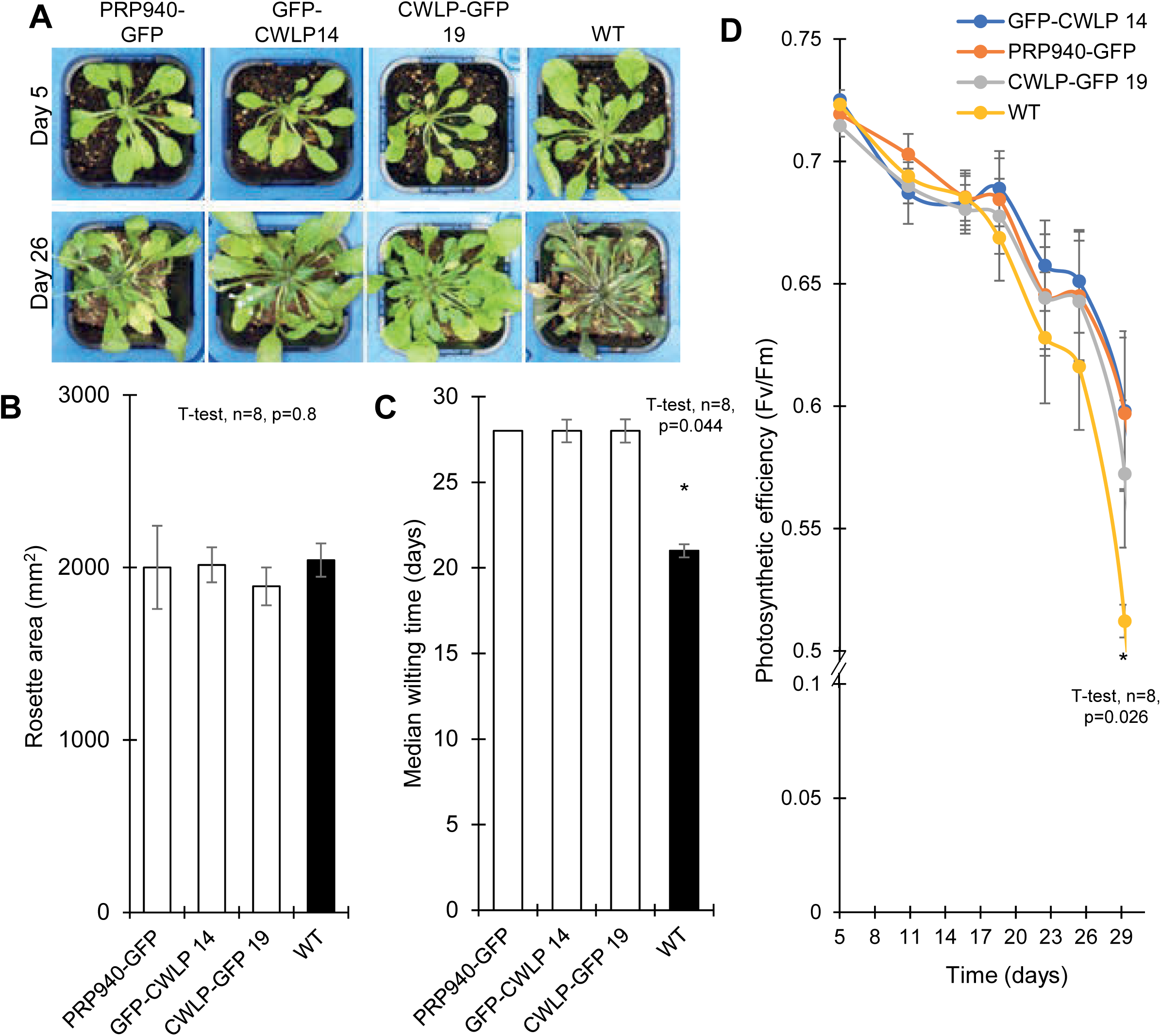
Overexpression of CWLP and PRP940 increases tolerance to water shortage. One-month-old PRP940-GFP, GFP-CWLP, CWLP-GFP, and WT plants were exposed to dehydration by water withholding. (A) PRP940-GFP, GFP-CWLP (line 14), CWLP-GFP (line 19), and WT plants, 5 and 26 days after the last irrigation (upper and bottom rows, respectively). (B) PRP940-GFP, GFP-CWLP, CWLP-GFP, and WT plants, 5 days after the last irrigation, had no significant difference in their rosette size (T-test, n=8, p=0.8). (C) During water withholding, wilting time of PRP940-GFP, GFP-CWLP, and CWLP-GFP plants is significantly longer compared with that of WT plants (T-test, n=8, p=0.044). (D) The photosynthetic efficiency (Fv/Fm) of WT plants decreases faster during water withholding than that of CWLP-GFP or PRP940-GFP plants (T-test, n=8, p=0.026). Fv/Fm was measured using PlantScreen – High-Throughput Phenotyping system.

Observed interactions of CWLP and PRP940 with aquaporins and CWLP-mediated recruitment of PP2A-B’’ regulatory subunits to PM clusters implied that CWLP (and most likely PRP940) not only anchors aquaporins to certain nanodomains but mediates their interactions with PP2A phosphatases. This option was also supported by earlier LC-MS/MS analysis of aquaporin cross-linked interacting proteins, which identified the abundant PP2A-A1 (RCN1, At1g25490) subunit in direct association with PIP1;2 and PIP2;1 (122). The formation of heterotrimeric complexes of A scaffold subunits with all five catalytic (C) and most regulatory (B, B’ and B’’) subunits has been recently confirmed by systematic mass spectrometry analysis of protein complexes purified from 13 different plant species, including Arabidopsis (131). To confirm aquaporin-association with CWLP-binding PP2A-B’’γ subunit, we co-expressed PP2A-B’’γ-YFP and PIP2;1-mCherry in *N. benthamiana* leaves and affinity purified PP2A-B’’γ-YFP protein complexes on GFP-Trap from the membrane protein fraction. Western blots probed with anti-mCherry monoclonal Ab and anti-PIP2;1 polyclonal Abs confirmed that the PP2A-B’’γ-YFP (detected by anti-GFP Ab, Fig 7E left panel) co-immunoprecipitated with aquaporin PIP2;1 and with other aquaporins, only detected by the polyclonal antibodies, that might be included in aquaporin heterotetramers (Fig. 7E, right panel). Ultimately, we co-expressed CWLP-CFP, PP2A-B’’γ-YFP, and PIP2;1-mCherry in *N. benthamiana* leaves (Fig. 7Fa-c, respectively). Co-localization of all three proteins in the PM was indicated by overlapping blue, yellow and red signals in merged images (Fig. 8Fd and fluorescence of numbered scanned trajectories shown in the middle panel). Similarly to the co-localization of CWLP-CFP and PP2A-B’’γ-YFP, the PM clusters containing all three proteins varied in size and included ball-shaped structures which protruded into the cytosol (Fig. 8F right panel). Whether these larger invaginated clusters correspond to more prominent microdomain bulges prepared for endocytosis by dynamin-mediated fission remains to be further studied. Recruitment of PP2A-B’’ subunits by CWLP suggested that PP2A enzymes might mediate dephosphorylation of CWLP-(and PRP940) interacting aquaporins, stimulating the closure of their water channels and thus reducing cellular dehydration. Our observation that CWLP and PRP940 overexpression confers remarkable tolerance to osmotic stress-induced plasmolysis in leaf cells also supported this hypothesis.

### 3.5 CWLP overexpression improves dehydration and freezing tolerance

To examine whether CWLP overexpression indeed improves plant dehydration tolerance, we first exposed three-month-old WT, and CWLP-GFP Arabidopsis plants, grown side by side under short-day (SD) conditions, to dehydration (Fig. S7). Ten days after stopping irrigation, 70% of the WT and only 20% of CWLP-GFP plants wilted. After 20 days, more than 90% of WT but only 65% of the CWLP-GFP plants were dehydrated (Fig. S7B and C). The relative water content of leaves at this time-point decreased to below 50% in WT plants while remaining above 60% in CWLP-GFP plants (Fig. S7D). To monitor the decline of photosynthetic activity caused by dehydration, one-month-old CWLP-GFP, GFP-CWLP, and PRP940-GFP Arabidopsis plants were grown in pots under SD photoperiod and subjected to dehydration (Fig. 8). After ceasing irrigation for 5 days all plants possessed similar rosette size (Fig. 8B). The WT plants wilted completely after 20 days, while PRP940-GFP, GFP-CWLP and CWLP-GFP lines showed delayed wilting, 27-28 days after water withdrawal (Fig. 8A and 8C). Monitoring photosystem II quantum yield (Fv/Fm) during the dehydration period indicated that the photosynthesis capacity started to decline faster in WT plants after 20 days of water withholding compared to the CWLP and PRP940 OE lines (Fig. 8D).

We also generated CWLP-GFP potato lines and tested their dehydration tolerance. Overexpression of CWLP-GFP was first confirmed by western blotting with anti-GFP Ab compared to Arabidopsis, and then PM-ND localization of CWLP-GFP was verified in potato leaf cells by confocal microscopy (Fig. S8A). As seen in Arabidopsis, the expression of CWLP-GFP in potato conferred delayed wilting compared to WT (Fig. S8B and 8C) and slowed down turgor loss of leaves (Fig. 8D).

Our survey of gene expression data (Fig. S1D) indicated that transcription of *CWLP* shows the highest induction in response to cold stress. To confirm the cold-inducibility of *CWLP*, WT Arabidopsis plants were exposed to 4°C in the light for four days and then transferred to 25°C for recovery. Although no cold-induced phenotypic damage was observed, we detected a gradual elevation of *CWLP* transcript levels during cold exposure. Compared to gradual increase of transcript level during the first 72 hours of cold acclimation, the recovery-induced decrease in *CWLP* transcription occurred fast, within 2 h, at 25°C (Fig. 9A). We compared freezing tolerance of WT, CWLP-GFP, N-del-CWLP-GFP, C-del-CWLP and *cwlp* (GABI_382D01, T-DNA K/O mutant, Fig. S9) lines by measuring the temperature at which 50% electrolyte leakage occurred (LT_50_) due to freezing-induced cellular damage (85). Detached leaves of 42-d-old non-acclimated (NA) and acclimated (ACC) plants, pre-treated for 14 days at 4°C, were exposed to a low-temperature range of -1°C to -22°C and then thawed under highly controlled conditions. Whereas non-acclimated plants of all lines were equally sensitive to freezing (Fig. 9B), the LT_50_ value of acclimated CWLP-GFP leaves was 1.5°C lower compared to acclimated WT (Fig. 9C). CWLP-GFP leaves showed the highest difference of LT_50_ value between NA and ACC plants (acclimation capacity) (Fig. 9C). Importantly, the *cwlp* mutant did not show an enhanced freezing sensitivity; and we could not observe phenotypic differences in comparison with WT and CWLP-GFP plants (Fig. S9B-D). This observation suggested that CWLP and PRP940 probably share redundant functions, which await confirmation by future analysis of double mutants.

**Figure 9.**
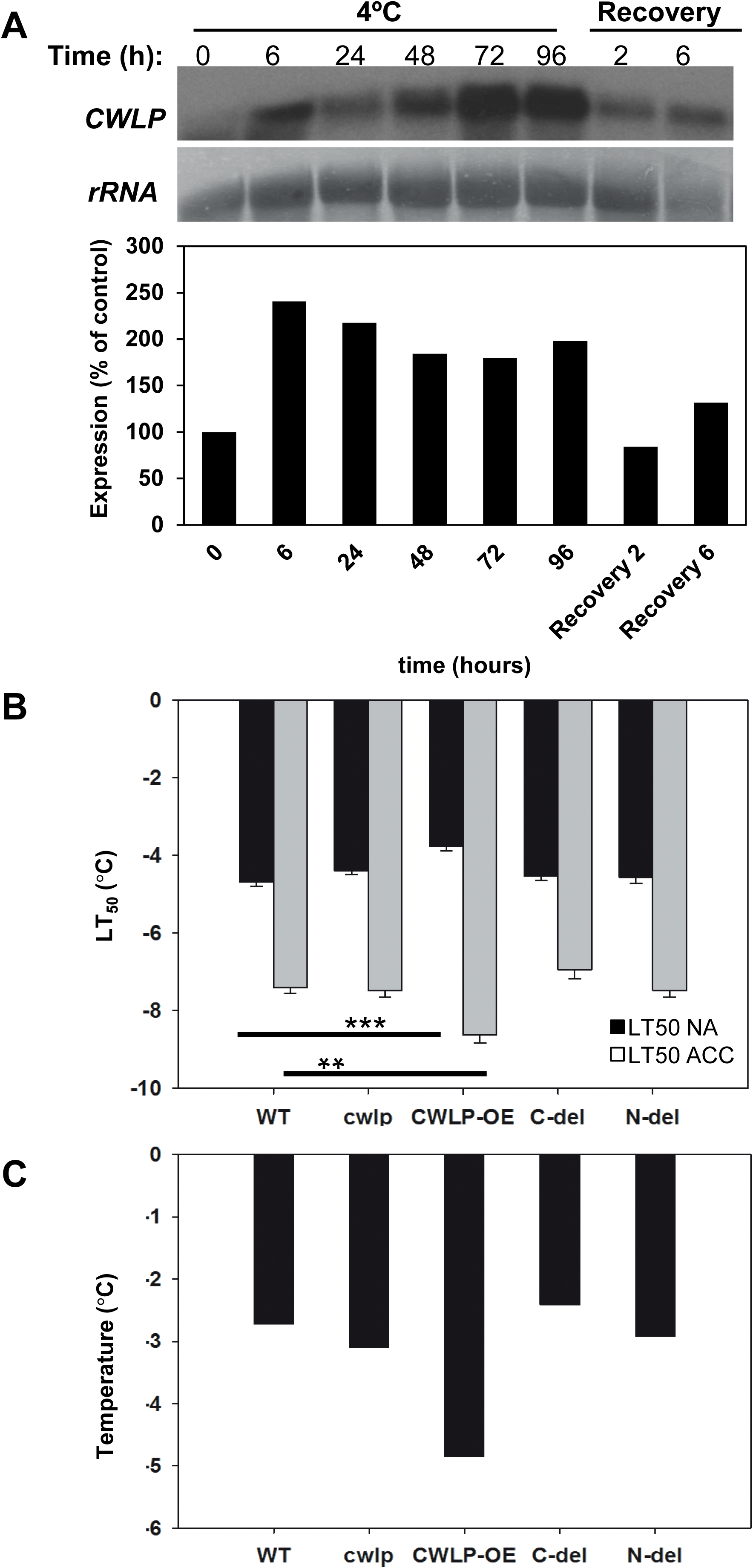
CWLP-GFP overexpression improves freezing tolerance. (A) Cold-induced gradual increase in *CWLP* transcription. Upper panel: Northern blot of total RNA extracted from leaves of 6-week-old WT plants during exposure to 4°C in the light for 96 h and 6h recovery at 25°C. The coding region of CWLP cDNA was used as a probe. Tomato DNA fragment of 26S rRNA was used as a probe for estimating RNA loading. Lower panel; ImageJ comparison of transcript levels relative to rRNA loading controls. (B) Comparison of electrolyte leakage (LT_50_) caused by gradually declining temperatures in detached leaves of five different lines: CWLP-GFP (denoted as CWLP-OE), WT (Col-0), *cwlp* mutant (GK382DO1), C-del-CWLP (denoted as C-del) and N-del-CWLP-GFP (denoted as N-del). Plants were exposed to increasing freezing stress from -1°C to -22°C with or without pre-acclimation at 4°C for 14 days, defined as acclimated –ACC (grey) or non-acclimated NA (black), respectively. SE of 3 leaves of 5 plants. Asterisks indicate significant difference compared with WT (**p <0.01, ***p <0.001) (C) Acclimation capacity deduced from (B), showing the minimal temperature after which, each acclimated line survived.

## 4 DISCUSSION

### 4.1 CW-PM linker functions of CWLP and PRP940

Our study provides an insight into previously proposed but yet uncharacterized functions of CWLP and PRP940 “cell-wall-membrane linker” proteins, which are Arabidopsis homologs of *Brassica napus* BnPRP, the first described member of C-type HyPRPs (1, 65). We found that cellular localization of N- and C-terminal GFP fusions of CWLP and PRP940 overlaps with the CW and PM. The N-terminus of CWLP is located in the CW vicinity, whereas C-terminal GFP-tags of both proteins are detected at the cytosolic surface of PM. We detected weak glycosylation of CWLP, which correlates with its anomalous behavior in SDS-PAGE (Fig. 6) and suggests that glycan sidechains of CWLP and PRP940 might anchor these proteins to yet unknown components of the CW. In fact, both proteins were identified in purified CW fractions (69), and potentially glycosylated hydroxyproline (HyP) residues were detected in PRD of PRP940 by mass spectrometry (50). To better understand their CW interactions, further study of the composition and interactions of glycosylated N-terminal PRDs is necessary.

CWLP and PRP940 display clustered co-localization in the PM with aquaporin PIP2;1 in nanodomains (103). Our study confirms that C-terminal 8CM domains of both proteins are closely related to 8CMs of Hydrophobic Protein from Soybean (PDB entry 1HYP; 94) and nsLTP proteins, which form tunnel-like lipid-binding cavities and might function as PM anchors. Although recent data suggest S-acylation of CWLP and PRP940 8CM Cys residues (97), little is known thus far about the identity of bound lipid residues and their roles in membrane trafficking and PM anchoring.

Lipid charging, S-acylation, and glycosylphosphatidylinositol (GPI)-anchoring of PM proteins promote their incorporation into detergent-resistant membranes (DRMs) of PM nanodomains (14, 132, 133). Our data demonstrate that CWLP and PRP940 are DRM-resident proteins localized in PM-NDs. While AZI1 and AZI1-like (AZL) small HyPRP proteins are implicated in non-vesicular lipid exchange in ring-like structures around the chloroplasts (64), we failed to identify CWLP and PRP940 in such structures (see, e.g., Fig. S5).

Cell wall linker function of CWLP is well-illustrated by the overexpression of CWLP which counteracts hyperosmosis-stimulated plasmolysis of epidermal cells. This suggests that upon its natural induction by cold stress, CWLP functions as a linker preventing destructive separation of CW from the PM, while the intrinsically disordered structure of its PRD (Fig. S1E) might provide flexible contact with the CW. Microsequencing data of DRM proteins demonstrated elevation of CWLP and PRP940 content during cold acclimation (134). It was accompanied by simultaneous increase of CW-relaxing enzymes including expansins, glycoside hydrolases (GHs), pectin methylesterases (PMEs) and xyloglucan endotransglucosylase/hydrolases (XTHs), capable of reducing cellular damages caused by ice nucleation at subzero temperatures (135). Removal of the N-terminal part of PRD or the last eight amino acids of 8CM abolishes the ability of CWLP to prevent plasmolysis, indicating that both termini are necessary for its linker function. CWLP and PRP940 thus appear to function like “pickets” linking the PM “curtain” to the CW contributing to its attachment when decreased cellular turgor leads to protoplast shrinkage. The assembly of such “pickets” appears to be supported by the cortical actin filament network (136). Thus, enzymatic removal of the CW during protoplast preparation does not cause a significant alteration in the positioning of CWLP-marked PM-MD clusters, similarly to several other DRM proteins (137). Nanodomain localization of CWLP (and PRP940) is also unaffected by disruption of the microtubule network in response to oryzalin treatment. By contrast, inhibition of actin polymerization by LatB, which still allows initiation of early endosomes carrying earlier-formed CWLP clusters, blocks their further endocytotic cycling. Correlatively, inhibition of actomyosin network-dependent trafficking by a mutated form of myosin XI-K tail (138) results in even distribution of CWLP in the PM.

Internalization of nanodomain proteins, such as aquaporins (98) through both CME and CIE pathways (10) requires dynamin-mediated fission of PM invaginations to release endocytic vesicles (29). Our data support the observation that dynamin-aided endosome formation occurs in plants despite LatB-mediated inhibition of actin polymerization (25, 40). Moreover, our mass spectrometry data showed that CWLP is directly associated with Dynamin-Related Proteins DRP1A and DRP1C, which likely govern its membrane trafficking (139). The association of CWLP with DRP1s detected in the cytoplasmic fraction supports the notion that dynamins form cytoplasmic complexes (e.g., in the ER) with diverse adaptor proteins, in order to target them to various subcellular locations (140–142). Although DRP1A and DRP1C interaction with CWLP does not indicate their dimerization and oligomerization, required for GTPase function and fission of PM invaginations (143–145), still mutation of DRP1A compromises endocytosis (118, 146). Additionally, transcription of functionally redundant *DRP1B* and *DRP1E* genes is highly induced during cold acclimation, similarly to the cold-induced accumulation of CWLP (22, 134), while DRP1E mutagenesis confers reduced freezing tolerance (147).

### 4.2 Recruitment of aquaporins and PP2A phosphatases by CWLP and PRP940: a model for the closing aquaporin water channels and reducing water loss

The observations that CWLP and PRP940 overexpression led to enhanced whole-plant tolerance to water shortage further strengthened the contribution of overexpressed CWLP and PRP940 to overcome water stress. This dehydration withstanding ability was also confirmed in transgenic potato overexpressing CWLP-GFP, which showed similar CWLP-GFP pattern of PM clustering, demonstrating that tolerance to water shortage is not specific to Arabidopsis and might be implied to other plants.

Exposure to freezing temperatures induces similar cellular changes as dehydration stresses because the formed ice crystals reduce water potential in the apoplast and extracellular space, promoting water sequestration from cells. During this process, called equilibrium freezing-induced dehydration, the CW is remodeled to be more elastic for coping with the physical stress of ice crystallization, including the synthesis of protecting wax, cutin, or suberin layers. Severe dehydration may lead to lethal injury by cytorrhysis (i.e., irreversible plasmolysis), separating the deformed CW from the PM. This damage can be reduced by: i) increasing CW flexibility; ii) elevating the density of CW-PM linker molecules, and iii) limiting cellular water loss by closing aquaporin water channels in the PM (135, 148). Co-localization, split Y2H interaction, and co-immunoprecipitation of aquaporins PIP2;1 and PIP2;7 with CWLP and PRP940 suggest that they might also modulate the activity of aquaporins. The finding that flotillin FLOT2, which also interacts with aquaporins (43), did not co-immunoprecipitate with CWLP or PRP940 indicates that CWLP and PRP940 mark a novel type of distinct nanodomain complexes. Moreover, the demonstration that CWLP interacts in Y2H assays with several PP2A-B’’ subunits and interact *in vivo* with PP2A-B’’γ subunit and PIP2;1, and all the three are co-localised, implies that CWLP and likely PRP940 recruits PP2A phosphatases to aquaporins. Notably, we found that the bulk of cytoplasmic PP2A-B’’γ subunit is targeted to PM-NDs by transient co-expression of CWLP.

Compared to intensive phosphoproteomic and structural studies of aquaporin phosphorylation (149–152), less is known about protein kinases involved in regulation of aquaporins’ activities. Aquaporins were reported to undergo phosphorylation on conserved Ser residues in their cytoplasmic B-loops and C-terminal domains. Nonetheless, it is still debated which N- or C-terminal phosphorylation sites mediate the conformational change required for opening the water channel (153, 154). Spinach aquaporin PM28A is phosphorylated on Ser115 and Ser274 residues by Ca^2+^-dependent kinases, which is prevented by okadaic acid, an inhibitor of PP1 and PP2A phosphatases (155). Arabidopsis PIP2;1–2; 4 and PIP2;7 undergo phosphorylation on the C-terminal Ser 283 and Ser286 residues by the transmembrane Sucrose-Induced Receptor Kinase (SIRK1) and on Ser280/Ser283, stimulating gate opening of aquaporins in response to ethylene signaling (156, 157). Phosphorylation of Ser283 is also required for PIP2;1 trafficking from the ER to the PM (158–161).

On the other hand, PP2A was reported to be involved in dephosphorylation of PM aquaporins causing low-temperature-induced closure of tulip petals (162, 163). Furthermore, phosphorylation status of Ser_35_, Ser_116_ and Ser_274_ in the tulip aquaporin could influence the activity of its water channels (164). The Ser_35_ residue is highly conserved in 12 of the 13 Arabidopsis PIPs (except for AtPIP2;4). Additionally, the opening and closure of the spinach aquaporin PM28A (PIP2;1) was found to be regulated by phosphorylation/dephosphorylation, respectively, of Ser_115_ in the first cytoplasmic loop (B loop), and Ser_274_ in the C-terminal part (153). Thus, collectively these findings support our suggested model (Fig. 10), in which CWLP and likely PRP940 form PM-ND complexes with aquaporins that recruit PP2A phosphatase via its B’’ regulatory subunits and promote aquaporin cytoplasmic gate closure by dephosphorylation.

**Figure 10.**
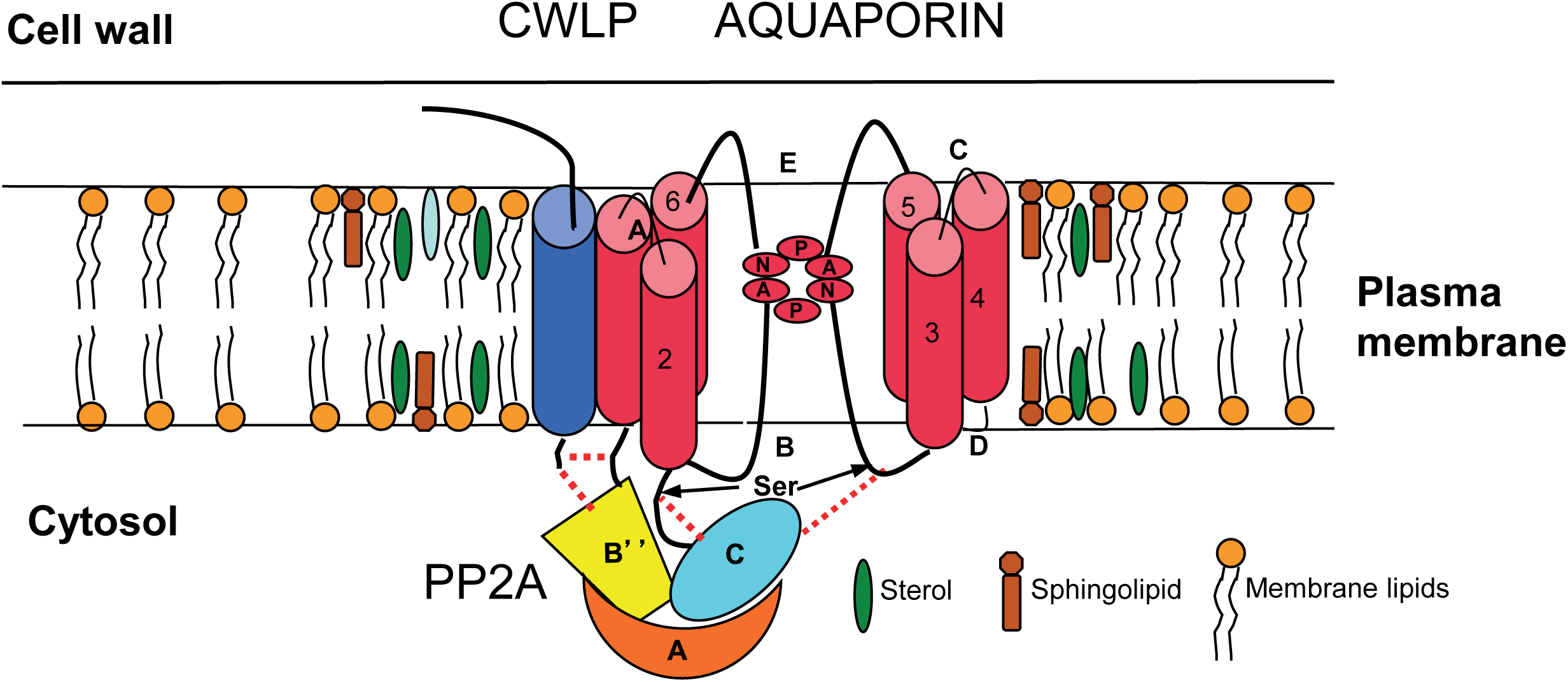
A model suggesting the involvement of CWLP in targeting PP2A to aquaporins in nanodomains according to protein-protein interactions. CWLP (blue), with N-terminus anchored to the CW and C-terminus protruding to the cytosol, serves as a nanodomain scaffold, which interacts with aquaporins in PM-NDs. A single aquaporin monomer out of the tetrameric complex is shown with enlarged B and E loops, which place the dual asparagine-proline-alanine (NPA) motif in the center of the water channel. Both CWLP and aquaporins interact with PP2A-B” regulatory subunits, facilitating the recruitment of trimeric PP2A enzyme (A, B” and C subunits) to the vicinity of cytoplasm-facing Ser residues of aquaporin loops, likely mediating their dephosphorylation and thereby the closure of the cytoplasmic water gate. Increased incorporation of CWLP molecules into the PM (i.e., under cold acclimation or overexpression) enhances aquaporin closure by preventing the decrease of cellular water potential. The CWLP homolog PRP940 is predicted to function similarly. molecules into the PM (i.e., under cold acclimation) enhances aquaporin closure by preventing the decrease of cellular water potential. The CWLP homolog PRP940 is predicted to function similarly.

Among the different types of nanodomains (165), so far, little is known about the potential association of aquaporins with remorin- and formin-containing PM clusters. By contrast, aquaporins were identified in association with Arabidopsis FLOT2 by mass spectrometry (43) and in mammalian flotillin complexes (166). It is still unknown how Arabidopsis flotillins recruit aquaporins and influence their activities, while studies of human flotillins show their involvement in Epidermal Growth Factor Receptor tyrosine kinase signaling in many kinds of human cancers (167), and their phosphorylation by PKC (Protein Kinase C) at Ser315, which stimulates the internalization of flotillin and associated cargos. Dephosphorylation of this residue by PP2A stabilizes nanodomain localization of FLOT1, enhancing cell migration and formation of endothelial barriers (44, 46, 168). Such studies indicate that various protein kinases and PP2As might be involved in modulating activities, interactions, and localization of nanodomain scaffold proteins. Thus, exploring PP2A-mediated dephosphorylation of aquaporins and PP2A interaction with CWLP and PRP940 may uncover many yet unknow aspects in the regulation of nanodomain functions and dynamics.

From the viewpoint of aquaporin regulation, it is intriguing that all CWLP-interacting PP2A-B’’ subunits contain EF-hand motifs similar to those of calmodulins (CaMs). Whereas the 8CM domain of Arabidopsis snLTP1 is found to bind calmodulin (CaM) in a Ca^2+^-dependent manner (169), CaM-binding to human aquaporins inhibits their activity, which is counteracted by phosphorylation of C-terminal Ser residues (170). Therefore, additional studies of DRM complexes are required to determine whether calmodulins, other cargos, protein kinases, known scaffold proteins, signaling factors, and AF network components are associated with CWLP- and PRP940-marked nanodomains.

CWLP overexpression, confers enhanced dehydration tolerance in both Arabidopsis and potato plants. These observations suggest that in addition to counteracting plasmolysis by its CW-PM linker function, CWLP probably enhances the closure of aquaporins’ gates in all cells where it is expressed. In WT Arabidopsis plants transcription of *CWLP* increases steadily during cold acclimation but declines fast when plants are returned to normal temperature. This elevation of *CWLP*, and to lesser extent, *PRP940*, is accompanied by co-expression of many other genes involved in protective responses to cold and freezing tolerance (134). Cold acclimation is thus essential for natural accumulation of CWLP in the CW-PM region of cells and formation of PP2A-interacting CWLP-aquaporin complexes in nanodomains. Further characterization of assembly and dynamics of the initially defined tri-partite regulatory complexes formed in PM-ND based on provided CWLP or PRP940 scaffold as well as the stress-inducible expression of their components may therefore provide valuable tools for practical applications aiming at the improvement of freezing and dehydration stress tolerance.

## Supplemental Information

The following data is available online:

**Figure S1**. CWLP and PRP940 expression, cellular localization, and topology.

**Figure S2**. ConSurf analysis of evolutionary conservation of AtCWLP 8CM domain and AtPIP2;7.

**Figure S3**. PRP940-GFP is not co-localized with microtubules.

**Figure S4**. The effect of LatB on PM-localization patterns of CWLP-GFP and PIP2;1-mCherry in ectopically expressed in Arabidopsis plants.

**Figure S5**. Clustering of CWLP-GFP in the PM is not affected in CW-free protoplasts prepared from leaves of CWLP-GFP expressing Arabidopsis plants.

**Figure S6**. PP2A-B”γ-YFP is not co-localized with Golgi and endosomal markers upon transient expression in *N. benthamiana* leaves.

**Figure S7**. CWLP overexpression increases tolerance to water shortage.

**Figure S8**. Overexpression of CWLP-GFP in potato confers improved dehydration tolerance.

**Figure S9**. Phenotype of *cwlp* T-DNA knockout line does not differ from that of WT.

**Figure S10**. AtCWLP constructs used in this study.

**Table S1**. Peptides detected by LC-MS/MS analysis of bands isolated from the gel shown in Figure 5A.

**Table S2**. Oligonucleotide primers used in this study.

**Table S3**. Antibodies used in this study.

**Supplemental Methods**

## Acknowledgments

We thank Sebastian Bednarek for anti-DRP1 antibodies, Nadav Sorek for the advice on extracting DRMs, and Shaul Yalovsky for the GFP-ROP6^CA^ probe. We are grateful to Horst Röhrig and Dorothea Bartels for the immunoprecipitation method. This work was supported by BARD, The United States – Israel Binational Agricultural Research and Development grant (IS-4143-08 R) to A. Z., B. L., and E. S., and DFG, German – Israeli – Palestinian Trilateral grant (KO-1438/13-2) to A. Z and C. K.

## AUTHORS’ CONTRIBUTIONS

A. Z., Y. F., A. H., H. E., E. S., B. L., Y.J. L and C.K. designed the research. A. H., H. E., Y. F., T. H., D. A., G. E., N. R., H. S, E. B., O. E, S. S, and M.A. A. performed the experiments. E. Z. designed and carried out the freezing tolerance experiments. Results were discussed with all authors. The article was written by A.Z., C. K, Y. F., H. E., A. H., B. L., and E. S., read by all authors.

## Notes

### Competing Interest Statement

The authors have declared no competing interest.

