## Supporting information for "CWLP and PRP940 form plasma-membrane nanodomain complexes with aquaporins, interact with PP2A and contribute to dehydration tolerance"

#### Table S1

Peptides detected by LC-MS/MS analysis of bands isolated from the gel shown in Figure 5A.

| Sample# | Gene ID | Gene Name | # peptides | Peptide sequence |
|---|---|---|---|---|
| CWLP-GFP IP | At5g42080 | Dynamin 1A | 60 | AAQSEIDAVAWSK |
| | | | | ACTALGDHGDSSALPTLWDSLPAIAVVGGQSSGK |
| | | | | AGGEKVYNVFDNQLPAALKR |
| | | | | DDGTREYAEFLHLPR |
| | | | | EREYFSNTTEYR |
| | | | | EVDPSGDRTFGVLTK |
| | | | | EYAEFLHLPR |
| | | | | EYFSNTTEYR |
| | | | | FTDFAAVR |
| | | | | FTDFAAVRK |
| | | | | GGNPTHSIFDR |
| | | | | GGNPTHSIFDRYNDSYLR |
| | | | | GTDAVEILEGR |
| | | | | GYQPHLIAPEQGYR |
| | | | | IDDGTREYAEFLHLPR |
| | | | | IDLMDKGTDAVEILEGR |
| | | | | IILAISPANQDLATSDAIK |
| | | | | IPGIQSLINK |
| | | | | KFTDFAAVR |
| | | | | KKFTDFAAVR |
| | | | | KLVTEADGYQPHLIAPEQGYR |
| | | | | KLVTEADGYQPHLIAPEQGYRR |
| | | | | LFDQIFKEHLDGVR |
| | | | | LGKPIAADAGGK |
| | | | | LIESSIVSIR |
| | | | | LKYPWVGVVNR |
| | | | | LPGLTKVAVDGQSDSIVK |
| | | | | LQFDKQLAMDNIR |
| | | | | LQFDKQLAMDNIRK |
| | | | | LSSLLNEDPAIMER |
| | | | | LVTEADGYQPHLIAPEQGYR |
| | | | | LVTEADGYQPHLIAPEQGYRR |
| | | | | LYSIMEICR |
| | | | | MENLISLVNK |
| | | | | MENLISLVNKIQR |
| | | | | NKNVDMIAAR |
| | | | | QDVEKGGNPTHSIFDR |
| | | | | QLAMDNIR |
| | | | | QLAMDNIRK |
| | | | | QLAMDNIRKLVTEADGYQPHLIAPEQGYR |
| | | | | RLSSLLNEDPAIMER |
| | | | | RPLVLQLQK |
| | | | | SIVYCQVR |
| | | | | SLLDHFFAELGTMDMKR |
| | | | | SQADINKNVDMIAAR |
| | | | | SRIPGIQSLINK |
| | | | | SSVLESIVGK |
| | | | | SSVLESIVGKDFLPR |
| | | | | SVNETVELK |
| | | | | SVNETVELKQYPALR |
| | | | | SYIEKPNCIILAISPANQDLATSDAIK |
| | | | | TVLELETELSR |
| | | | | VAVDGQSDSIVK |
| | | | | VAVDGQSDSIVKDIENMVR |
| | | | | VEVTNAAIESLDK |
| | | | | VEVTNAAIESLDKMR |
| | | | | VEVTNAAIESLDKMREGSK |
| | | | | VYNVFDNQLPAALK |
| | | | | VYNVFDNQLPAALKR |
| | | | | YPWVGVVNR |

| Sample# | Gene ID | Gene Name | # peptides | Peptide sequence |
|---|---|---|---|---|
| CWLP-GFP IP | At1g14830 | Dynamin 1C | 49 | ACTVLGDHGGEGMSLWEALPTVAVVGGQSSGK |
| | | | | AVVYCQVR |
| | | | | DDIDAVAWK |
| | | | | EIEDETDRITGK |
| | | | | EKEKLGAMLDEDPQLMER |
| | | | | EKLGAMLDEDPQLMER |
| | | | | EQEYFETSPEYGHLASR |
| | | | | ESIVQDIENMVR |
| | | | | FADFAAVR |
| | | | | FADFAAVRK |
| | | | | FPTLASDIAAAANEALER |
| | | | | GPAEATVDAVHFVLK |
| | | | | IGRPIAVDSGAQLYTILELCR |
| | | | | IGSNVSAYINMVCDTLR |
| | | | | IPSIVALINK |
| | | | | IYGVFDHQLPAALK |
| | | | | KEIEDETDRITGK |
| | | | | KLHLEPEKEKPNPR |
| | | | | KSISETEELKR |
| | | | | KSISETEELKRFPTLASDIAAAANEALER |
| | | | | LDIMDKGTDCLDVLEGR |
| | | | | LGAMLDEDPQLMER |
| | | | | LGAMLDEDPQLMERR |
| | | | | LIDGSISYFK |
| | | | | LLSQHLETVIR |
| | | | | LQHPWVGIVNR |
| | | | | LVDMESSYLTVEFFR |
| | | | | MGSEYLAK |
| | | | | NAPAPNADPYSDNHFR |
| | | | | NAPAPNADPYSDNHFRK |
| | | | | QARDDIDAVAWK |
| | | | | QKIPSIVALINK |
| | | | | RKEQEYFETSPEYGHLASR |
| | | | | RVDMIAAR |
| | | | | SIDEINAELDR |
| | | | | SISETEELKR |
| | | | | SISETEELKRFPTLASDIAAAANEALER |
| | | | | SLIGLINK |
| | | | | SLLNFFYAQVGR |
| | | | | SSVLESVVGR |
| | | | | SSVLESVVGRDFLPR |
| | | | | SYVEKPNCIILAISPANQDIATSDAIK |
| | | | | TEDGTTEYAEFLHAPK |
| | | | | TFGVATK |
| | | | | TLIDLPGLTKVAVDGQPESIVQDIENMVR |
| | | | | VAVDGQPESIVQDIENMVR |
| | | | | VDMIAAR |
| | | | | VVSEADGYQPHLIAPEQGYR |
| | | | | VVSEADGYQPHLIAPEQGYRR |

**Table S2** Oligonucleotide primers used in this study.

| | 5' Primer | 3' Primer | **Amplified fragment size** | **Gene name and cloning description** |
|---|---|---|---|---|
| 1 | CTGCAGAAAAGGTAGACAAGTGTACGTC | TCTAGATTTTTGGAATGAGGGTTTGGG | 2000bp | CWLP putative promoter for GUS fusion. |
| 2 | GGTCGACATGGGGTCTCGCTCT | ACGTCGTACGGGTAAGCGGGACA | 1015bp | Full length CWLP for GFP fusion. |
| 3 | GGATCCATGGTGAGCAAGGGCGAG | TCTAGATTACTTGTACAGCTCGTCCATGC | 1014bp | mCherry for fusion to Full length CWLP. |
| 4 | GGTCGACATGGGGTCTCGCTCTCAAAA | GTCTAGAAGCGGGACATTTGAAGTCAG | 1016bp | Full length CWLP for CFP fusion. |
| 5 | GCGGCCGCTGCGACTGTACCCCTCCTAA | GCGGCCGCTTAAGCGGGACATTTGAAGTCA | 1018bp | Full length CWLP for N-Terminal GFP fusion |
| 6 | CTCGAGATGGGGTCTCGCTCTCAAAACCTCTCC | GAATTCTGGAGTTTTTCCACAGTCGAGAAGAAG | 978bp | C-terminal-Truncated CWLP (326a.a). |
| 7 | TGTACACCTCCGCCTACACCATGC | GGATCCAGCGGGACATTTGAAGTCA | 540bp | N-del-CWLP for GFP fusion (180 a.a). |
| 8 | GCATATGTGCGACTGTACCCCTCCTAAA | CTGCAGTTAAGCGGGACATTTGAAG | 940bp | Full length CWLP (w/o Signal Peptide) for Y2H nuclear system screen. |
| 9 | ATTAACAAGGCCATTACGGCCTGCGACTGTACCCCTCCTAAACCATCA | AACTGATTGGCCGAGGCGGCCCCAGCGGGACATTTGAAGTCAGATGG | 1021bp | Full length CWLP (w/o Signal Peptide) for Y2HSU screen. |

| | | | | |
|---|---|---|---|---|
| 10 | GTCTAGAATGGTGGATA CGGTTATTCCC | GTCTAGAAAAGGGAGCCT CCAGCGA | 1622bp | Full length PP2A-B” *(At5g28900)* for YFP fusion |
| 11 | GCGTCGACATGTTCAAG GTTGCAAGA | ACTCTAGACGCCTTGCTTA GAGTACCGATC | 1410bp | Full-length FLOT2 (*At5g25260*) for GFP fusion |
| 12 | ACGCCAAGGCAGAATGT TGT | CGGCTGCGTCTAGGTCAA GT | 57bp | *CWLP* qRT-PCR primers |
| 13 | TAGGTGCTTGCGTCGAC TTG | TTCGACTTCAACGAGGCCT T | 103bp | *PRP940* qRT-PCR primers |

**Table S3** Antibodies used in this study.

| Antibody | Manufacturer | Used dilution |
|---|---|---|
| Mouse anti-GFP | Covance | 1:1000 |
| Rabbit anti-GFP | Santa Cruz | 1:1000 |
| Rat anti-HA | Roche | 1:2000 |
| Rabbit anti-PIP2;7 | Agrisera | 1:2000 |
| Mouse anti-mCherry | Clontech | 1:1000 |
| Mouse anti-Tubulin . | | 1:100 |
| Rabbit anti-CWLP | This study | 1:10 |
| Cy3 goat anti-Rabbit IgG | Jackson ImmunoResearch Laboratories Inc. | 1:100 from 1mg/ml stock. |
| Alexa Fluor 488 goat anti-Mouse IgG | NTB | 1:100 from 2mg/ml stock. |

#### SUPPLEMENTAL METHODS

##### 1 Construction of CWLP expression cassettes and deletion mutants

***CWLP-HA*:** At*CWLP* coding sequence fused to a C-terminal HA tag was amplified using *Arabidopsis* cDNA as a template and primers (Table S2) and cloned between CaMV 35S promoter and NOS terminator and then inserted into the binary vector pPCV002 (Koncz *et al.* 1989).

***CWLP-GFP and PRP940-GFP*:** Translational fusion of CWLP-GFP driven by the 35S promoter was described in (Stein *et al.* 2011). Similarly, *PRP940* cDNA, C-terminally fused to GFP was cloned in pGIF as an expression cassette containing CaMV 35S promoter and OCS terminator (see Table S2 for primer details).

***pART27- CWLP-Δ326-334*:** Primers harboring *Xho*I and *EcoR*I sites (Table S2) were used to amplify the *CWLP* coding region (978bp) lacking the last 24 bp encoding PSDFKCPA. The CWLP-Δ326-334 fragment was inserted between CaMV 35S promoter and NOS terminator in the binary vector pART27 (Gleave 1992).

***Δ25-158-CWLP***: The N-terminal deleted CWLP was obtained by first cloning two complementary synthetic oligonucleotides encoding the signal peptide and harboring 5' *Cla*I and 3' *Bsp1407*I ends in the corresponding sites of pTZ57R (Thermo Scientific) to create pTZ57R-SP. Then, the *CWLP* coding region from nucleotide 475 to 1002 with 5'*Bsp1407*I and 3' *Bam*HI sites was cloned in frame in pTZ57R-SP using similar sites. The *Cla*I and *Bam*HI fragment was shifted to pGFPN and then the 35Spro-SP-N-delCWLP-GFP-OCSter cassette was cloned into a *Not*I site created in the polylinker of pPCV002.

***PP2A-B"γ-YFP***: Full-length cDNA of the PP2A-*B"γ* gene was synthesized (1608 bp) using RNA as template and cloned in the *XbaI* site of pBINPLUS-YFP (van Engelen *et al.* 1995).

***FLOT2-GFP*:** FLOT2 cDNA (S81324, obtained from ABRC) starting from the translation start site and lacking the last two codons (1410 bp) was fused in frame to GFP.

***CWLP Promoter-GUS*:** *Arabidopsis* genomic DNA (200 ng) served as a template for PCR amplification of 2 kb *CWLP* promoter region, including the 5'-UTR (25 bp) by specific primers (Table S2). After cloning in pTZ57R, the promoter was shifted as *Pst*I and *Xba*I fragment into pUC57 containing the *GUS* coding sequence with 189 bp tomato STLS-1 intron 2 and NOS terminator (Miller *et al.*, 2005). Then, the promoter-GUS cassette was cloned in the *Hind*III site of pGIF2 (Strizhov *et al.* 1996).

**2 Freeze Shattering and Immunolocalization**

Leaves were immersed in PME buffer (50 mM PIPES pH 6.9, 5mM EGTA, 2mM $MgSO_4$), 0.25% DMSO, 0.05% Triton X-100 and 1mM PMSF (Sigma #P7626) for 2 h, and then washed in PBS (137 mM NaCl; 2.7 mM KCl; 10 mM $Na_2HPO_4$; 2 mM $KH_2PO4$, pH7.4) 3 times for 5 min. Leaves were then freeze-shattered in liquid nitrogen (Wasteneys et al., 1997), incubated in PBS, 1% Triton X-100 for 1.5 h, and thereafter in PBS, 0.05% Triton X-100, 1% BSA overnight at 4°C. Two-hour incubation with rabbit anti-CWLP (1:10) or mouse anti-tubulin (1:100) (DM1A Sigma #T6199) in PBS followed. After three washes in PBS for 20 min each, leaf pieces were transferred either to Cy3-goat anti-rabbit IgG (1:100 from 1mg/ml stock, Jackson ImmunoResearch #115-165-072) or Alexa Fluor 488 goat anti-mouse IgG (1:100 from 2mg/ml stock, Molecular Probes #A-11029) in PBS and incubated for 2 h at 25°C. After three washes in PBS for 20 min each, leaf pieces were mounted in 0.01M phosphate buffer, 33% glycerol, 17% Mowiol (CM82070, Hoechst) on microscope slides and covered with cover-slips.

**3 Transient Expression in *N. benthamiana* Leaves**

Transient expression was performed using *Agrobacterium* GV3101 (pMP90RK) expressing the desired construct cloned in a binary vector (pPCV002, Koncz & Schell 1986), or pBINPLUS, (van Engelen *et al.* 1995). Cultures grown in LB medium overnight, were diluted with induction medium (50 mM MES [pH 5.6], 0.5% (w/v) glucose, 1.7 mM $NaH_2PO_4$, 20 mM $NH_4Cl$, 1.2 mM $MgSO_4$, 2 mM KCl, 17 µM $FeSO_4$, 70 µM $CaCl_2$, and 200 µM acetosyringone), and grown for additional 3-6 h until culture density reached $A_{600}$ 0.4. The two different *Agrobacterium* cultures were mixed, and the mixture was infiltrated into the leaves of 7-8-week-old *Nicotiana benthamiana* plants. The injected areas were inspected after 48 h by the confocal microscope or used for co-immunoprecipitation assays.

**4 Immunoprecipitation Using Whole Cell Extracts**

Leaf extract (in 50 mM phosphate buffer, pH 7.2, 150 mM NaCl, 2 mM $MgCl_2$, Complete Protease Inhibitors Cocktail (Roche), 2 mM PMSF, 140 mM β-mercaptoethanol) was incubated overnight with 4-6 µg rabbit anti-GFP polyclonal antibody (Santa Cruz) per ml of cell extract, with gentle shaking at 4ºC. The complexes were then adsorbed to 30 µl protein A Sepharose$^{TM}$ (Amersham Biosciences) for 3 h, washed twice with ice-cold wash buffer 1 (50 mM Tris-HCl [pH 7.5], 1 mM EDTA, 150 mM NaCl, 0.1% Nonidet P40), twice with wash buffer 2 (50 mM Tris-HCl [pH 7.5], 1 mM EDTA, 500 mM NaCl, 0.1% Nonidet P40) and once with wash buffer 3 (10 mM Tris-HCl [pH 7.5] and 0.1% Nonidet P40). Between each wash, the re-suspended beads were incubated for 20 min at 4ºC on a rotary shaker and centrifuged for 1 min at

12,000 g at 4°C. After the last wash, the proteins were eluted from the beads with SDS-PAGE sample buffer by boiling for 5 min (if not otherwise stated).

**5 Detection of CWLP glycosylation moieties**

CWLP-GFP expressing seedlings (700 mg) were ground in liquid nitrogen to a fine powder and, after adding 600 µl lysis buffer (50 mM Tris-HCl, pH 8.0; 1% SDS; 1 mM EDTA), were incubated at 100ºC for 5 min. The extract was diluted 1:10 with ice-cold wash buffer 1 (50 mM Tris-HCl, pH 7.5; 1 mM EDTA; 150 mM NaCl; 0.1% Nonidet P40) and centrifuged for 10 min at 16,000 g at 4ºC. The clear supernatant was collected and the CWLP-GFP was immunoprecipitated as described in section 4. After the last wash, the conjugated proteins were released from the Protein A-Sepharose-beads by boiling in 120 µl x2 Laemmli sample buffer and analyzed by SDS-PAGE using 10% acrylamide gels (115 µl were used for detection of glycosylation and 5 µl for western blotting). After protein blotting onto nitrocellulose membrane, one part of the membrane was stained with Glycoprotein Detection Kit (Sigma-Aldrich). The other part was probed with mouse anti-GFP monoclonal antibody (Covance), followed by detection with EZ-ECL (Biological Industries, Beth Haemek, Israel) using horseradish peroxidase (HRP)-conjugated goat anti-mouse IgG (Jackson Laboratories).

**6 Differential Fractionation of Membrane Proteins: Isolation of TSM and DRMs**

*Arabidopsis* seedlings (15-d-old), expressing either CWLP-GFP or the nanodomain resident protein GFP-ROP6$^{CA}$, were subjected to differential detergent protein extraction according to (Sorek *et al.* 2017), with the following modifications: The extraction buffer consisted of 50 mM phosphate buffer [pH 7.5], 2 mM $MgCl_2$, 25 mM NaCl, 10% glycerol, 2 mM ß-mercaptoethanol, 1 mM phenylmethylsulfonyl fluoride (PMSF), leupeptin and pepstatin. The cytosolic supernatant fraction (Sup) was collected after 100,000 g centrifugation for 60 min at 4$^0$C. The pellet was solubilized in an equal volume of extraction buffer supplemented with 1% (v/v) Triton X-100 and was gently shaken on ice for 30 min. After centrifugation at 56,000 g for 60 min at 4$^0$C, the supernatant containing Triton X-100 soluble membrane fraction (TSM) was collected. The pellet (DRM fraction) was dissolved in equal volume of extraction buffer containing 1% (v/v) Triton X-100 and 0.5% (w/v) SDS. After 30 min incubation on ice with gentle shaking and 15 min centrifugation at 20,800 g, the supernatant comprising proteins solubilized by SDS from the detergent resistant membranes (DRM or microdomains) was collected. Equal volume from each fraction was size resolved on 10%gel-SDS-PAGE and analyzed by immunoblotting using mouse anti-GFP or anti-HA monoclonal antibodies and then detected by goat anti-mouse secondary antibody and ECL.

**7 Identification of CWLP interactors in Triton-X-100-soluble cytoplasmic fraction by immunoblotting and LS-MS/MS analysis**

Total proteins were extracted from 3-d-old WT and CWLP-GFP-overexpressing etiolated seedlings in 50 mM Tris HCl [pH 8.0], 150 mM NaCl and 1% Triton X-100. After 1 h incubation at 4°C, and centrifugation at 100,000 g, the supernatant was filtered through a 0.45 μm Millex HV filter (Millipore). CWLP-GFP protein interactors were immunoprecipitated using anti-GFP antibody-conjugated to magnetic beads (Miltenyi Biotec). Proteins eluted from the anti-GFP beads were resolved by 7.5% SDS-PAGE, stained with silver nitrate, or used for western analysis with monoclonal anti-GFP, anti-DRP1A/ADL1, and anti-DRP1C/ADL1C antibodies (Kang et al., 2003a; Kang et al., 2003b). The anti-α-tubulin DM1A monoclonal Ab (Sigma) and an anti-actin Ab (Andersland et al., 1994) were also used as control probes.

For LC-MS/MS, polypeptides co-purified with CWLP-GFP were separated by SDS-PAGE and stained with GelCode™ Blue Safe Protein Stain (Thermo Scientific). Specific bands were excised and digested with trypsin and then analyzed by liquid chromatography-tandem mass spectrometry (LC-MS/MS). In order to identify the detected peptides (SI Table S1), spectrum data were analyzed using X! Tandem (www.thegpm.org) and loaded into Scaffold 3.0 (Proteome Software Inc.). The cutoff settings parameters were: minimum 2 unique peptides, protein identification probability >95%, and peptide identification probability >80%.

**8 Co-immunoprecipitation using total membrane protein fraction**

*N. benthamiana* leaves (600 mg) transiently co-expressing pair combinations of CWLP-HA and GFP-PIP2;7, PP2A-B” γ-YFP with CWLP-HA, PP2A-B” γ-YFP and PIP2;1-mCherry, CWLP-HA and FLOT2-GFP, FLOT2-GFP and PIP2;1-mCherry or PRP940-GFP and PIP2;1-mCherry were ground in liquid nitrogen, homogenized in 1 ml extraction buffer (50mM Na-phosphate buffer [pH 7.2], 150 mM NaCl, 2 mM MgCl2, Complete Protease Inhibitor Cocktail (Roche), 2 mM PMSF, 140 mM β-mercaptoethanol) and centrifuged for 10 min at 20,800 g, at 4°C. The pellet was solubilized by homogenization in 500 μl cold extraction buffer, containing 0.5% (v/v) Triton X-100 and 0.5% SDS, was incubated on ice for 15 min with frequent gentle shaking, and then centrifuged at 20,800 g for 15 min. The pellet containing TSM and DRM fractions was resuspended in 1 ml wash buffer 1 (50 mM phosphate buffer [pH 7.5], 1 mM EDTA, 150 mM NaCl, and 0.1% (v/v) Nonidet P40). IP was carried out by adding anti-GFP polyclonal antibody crosslinked by DSP to protein A-agarose beads (Pierce), to half (0.5 ml) of the solubilized membrane fraction, and incubated overnight on a roller at 4°C. The other half (control) was similarly incubated but with protein A agarose with no crossed-linked antibody, serving as a control. Alternatively, GFP-Trap (Chromotek) was used for pull-down proteins interacting with GFP-PIP2;7, PRP940-GFP, FLOT2-GFP and PP2A-B” γ-YFP. Binding was performed according to the manufacturer protocol using membrane fraction (TSM and DRM). Following removal of 5 μl of input sample, the membrane fraction was 100-fold diluted and incubated on a roller at 4°C with 20 μl GFP-Trap for 4 hours. The bound protein complexes were washed as described in section 4, except for using phosphate-based buffer instead of Tris-HCl. Then, the protein complexes were eluted from the beads by 5 min boiling in 75 μl SDS-PAGE sample buffer, resolved on 7.5% SDS-PAGE, blotted onto nitrocellulose membranes, and separately probed with either mouse anti-GFP (BioLegend), rat anti-HA (Roche and BioLegend) monoclonal Abs or rabbit anti-PIP2;7 polyclonal Ab. The secondary antibodies used were HRP conjugated goat anti-mouse, anti-rat or anti rabbit IgG, respectively, and the antibody labelled proteins were detected by ECL.

**9 Yeast Two Hybrid (Y2H) Screens**

Nuclear interaction assay: This yeast-2-hybrid analysis was performed using the MATCHMAKER (Clontech) GAL4 two-hybrid system. The full-length coding region of CWLP, lacking its signal peptide (Fig. 1 E), was cloned in frame downstream to the GAL4 DNA binding domain in the yeast bait vector pGBKT7 (Clontech). This plasmid was co-introduced together with pACT2 vector (containing a GAL4 activation domain) harboring a cDNA expression library prepared from dark-grown *Arabidopsis* cell suspension (Salchert et al., 1998),into the yeast AH109 strain containing the GAL4 regulated marker genes *ADE2*, *HIS3* and *lacZ* (Clontech). These reporter genes are expressed when *in-vivo* two-hybrid interaction combines the GAL4 binding and activation domains (AD+BD). pACT2 clones capable of activating the *HIS3* and *lacZ* reporter genes were rescued, and their cDNA inserts were sequenced. *lacZ*, *ADE2* and *HIS3* activation was re-checked several times by "one on one" yeast-two-hybrid interaction for all the positive clones to eliminate false positive interactions.

Split-ubiquitin yeast-2-hybrid assay for membrane protein interaction: This analysis was performed using DUAL-HUNTER (DualSystems Biotech) yeast two-hybrid system. The full-length coding region of CWLP, lacking its signal peptide, was cloned in frame between the yeast *invertase* (*SUC2*) signal peptide and the C-Ub sequences in the pBT3-SUC bait vector (DualSystems Biotech). The fusion to the *invertase* signal peptide targets the CWLP to the plasma membrane in yeast cells with the C-Ub domain protruding towards the cytoplasm. This plasmid was co-introduced with a prey *Arabidopsis* cDNA library cloned in pPR3-N carrying the N-Ub domain (DualSystems Biotech Cat. #P02210) into the NMY51 strain containing the *ADE2*, *HIS3,* and *lacZ* reporter genes (DualSystems Biotech). These reporter genes are expressed when *in-vivo* two-hybrid interaction on the inner side of the plasma membrane combines the C-Ub and the N-Ub domains, thus releasing the VP16 transactivation factor, which in turn is capable of activating the *ADE2, HIS3,* and *lacZ* reporter genes in the yeast nucleus. pPR3-N clones capable of activating the *ADE2*, *HIS3,* and *lacZ* reporter genes were rescued, and their cDNA inserts were sequenced. *lacZ*, *ADE2,* and *HIS3* activation was re-checked several times by "one on one" yeast-two-hybrid interaction for all the positive clones to eliminate false positive interactions.

**10 Bioinformatic Analyses**

NCBI and TAIR databases were used for data mining and obtaining protein sequences and gene information (http://www.ncbi.nlm.nih.gov/, https://www.arabidopsis.org). Prediction of protein topology was performed using mobiDB (https://mobidb.bio.unipd.it.) for transmembrane domains, and PONDR for natural disordered regions (http://www.pondr.com/pondr-tut4.html.) Plant secretory signal peptides were identified by TargetP 2.0 (https://services.healthtech.dtu.dk/service.php?TargetP-2.0). Multiple alignment of protein sequences for cladograms of HyPRP proteins from *Arabidopsis* and other plants was performed by "ClustalW" software (www.ebi.ac.uk/clustalw), and MSA files were then used by "Mega4" software (http://www.megasoftware.net/) or by the DNAStar

Megalign Pro software, to obtain graphic presentations. The analyses of evolutionary-conserved residues and structure of the 8CM domain of HyPRPs and PIPs were carried out using the "Consurf" server (https://consurf.tau.ac.il/consurf_index.php) (Ashkenazy et al., 2010). The Bar (http://bar.utoronto.ca) and Genevestigator (https://genevestigator.com) servers were used for collecting gene expression data on the effects of abiotic stresses. Analysis of *CWLP* promoter was performed by using "plantCARE", (http://bioinformatics.psb.ugent.be/webtools/plantcare/html).

### SUPPLEMENTAL FIGURES

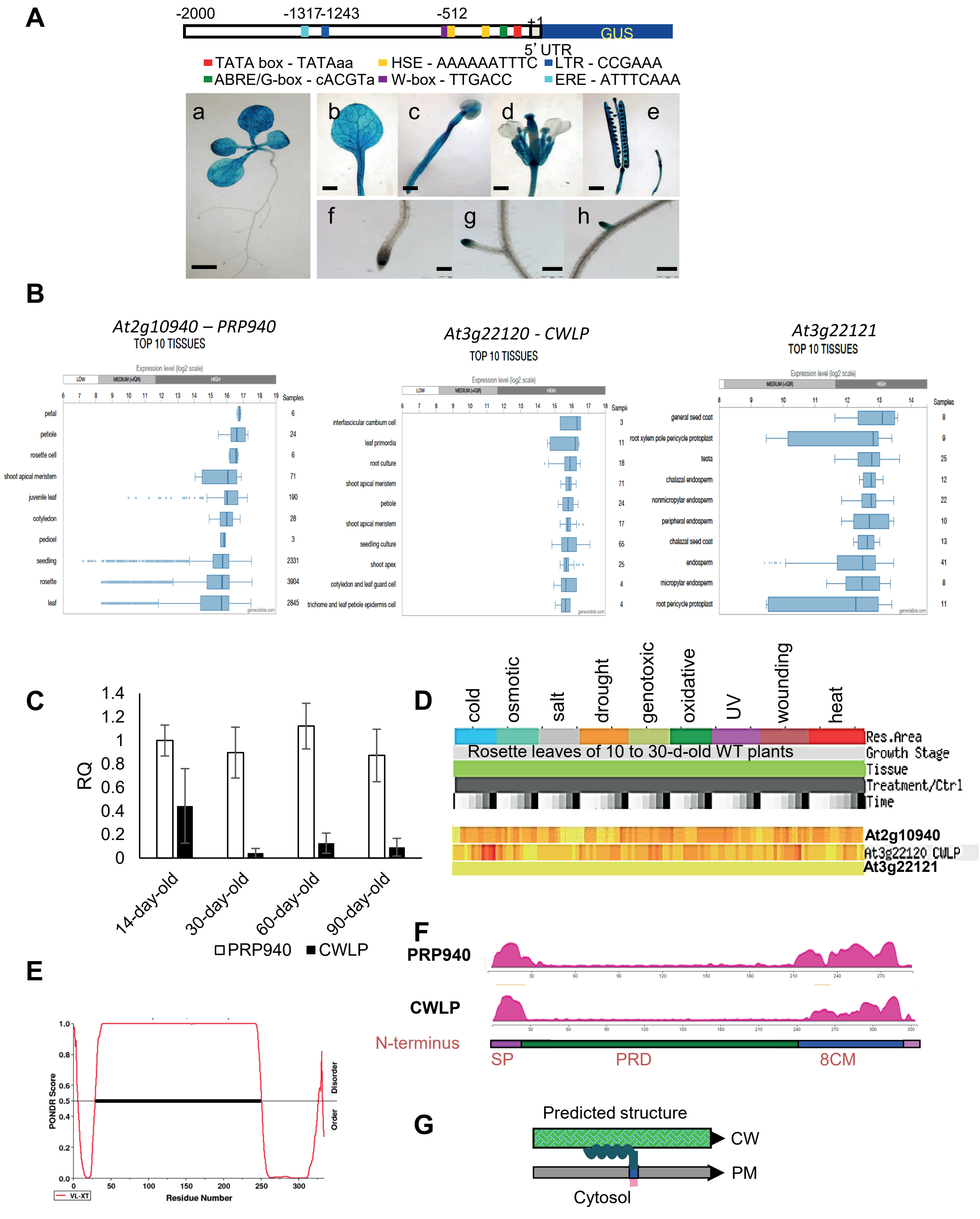


**Figure S1**
CWLP and PRP940 expression, cellular localization, and topology.
(A) CWLP promoter activity monitored by histological staining β-glucuronidase in pCWLP-GUS plants. (a) Seven-d-old seedling; (b) Mature rosette leaf; (c) Stamen; (d) Flower; (e) Silique; (f) Main root tip; (g) and (h) Lateral root tips of 10-d-old seedling. Bars in (a) 1mm, and in (b) to (h) 0.1mm. Schematic presentation of CWLP promoter indicates the positions of putative stress-related transcription factor binding sites ABRE (abscisic acid response element), HSE (heat stress responsive element), and LTR (low temperature responsive element) identified by the "plantCARE" (http://bioinformatics. Psb.ugent.be/ webtools/plantcare/html) search tool.
(B) Transcript levels of *PRP940* (*At2g10940*), *CWLP* (*At3g22120*), *lncRNA* (*At3g22121*, antisense of *CWLP*) during various stages of development, detected by using Affymetrix ATH1 arrays representing 111 tissues. Ten tissues exhibiting the highest expression values are presented:
(https://genevisible.com/tissues/AT/Ensembl%20Gene/AT2G10940;
https://genevisible.com/tissues/AT/Ensembl%20Gene/AT3G22120;
https://genevisible.com/tissues/AT/Ensembl Gene/AT3G22121)
(C) Comparison of *PRP940* and *CWLP* transcript levels in rosette leaves of 14, 30, 60, and 90-day-old plants grown in SD light period by qRT-PCR. Reactions were performed with 3 biological repeats, each with 2 technical replicates. RQ values refer to relative transcript levels compared with *PRP940* expression on the 14$^{th}$ day. Cyclophilin (*At2g36130*) transcript levels served as constitutively expressed control.
(D) Summary of transcript levels of *CWLP* (*At3g22120*), *lncRNA* (*At3g22121*) and *PRP940* (*At2g10940*) during exposure to different abiotic stresses, obtained by the Expression Angler software at the Bio-Analytic Resource (BAR) site (http://bar.utoronto.ca). The data was provided by Kilian et al. (2007) Plant J. 50: 347-363. All imposed stresses were measured within 24 h after the beginning of the following treatments: cold – continuous 4° C, osmotic (300 mM mannitol), salt (150 mM NaCl), drought (15 min exposure to air stream, reaching approximately 10% fresh weight loss), genotoxic (1.5 μg/ml bleomycin, 22 μg/ml mitomycin), oxidative (10 μM methyl viologen), UV (15 min UVB light), wounding (punctuation of leaves with 16-needle-pin tool), heat (3 h at 38$^0$C followed by recovery at 25$^0$C) stress treatments. (E) Prediction of Natural Disordered Regions (PONDR) in CWLP amino acid sequence (http://www.pondr.com/pondr-tut4.html), showing that the PRP domain is highly disordered, whereas the 8CM domain is well ordered. Similar prediction was obtained for PRP940. (F) Comparison of CWLP and PRP940 topology: Prediction of trans-membrane domains performed by using mobiDB prediction at https://mobidb.bio.unipd.it for Q9LIE9 (CWLP) and Q9SKI0 (PRP940). The 8-CM domain contains three predicted TM motifs, while the TM motif in the N-terminal secretory signal peptide is cleaved during the processing in the ER. (G) Suggested membrane topology of CWLP with N-terminal end in the CW vicinity and C-terminus protruding with few last amino acids into the cytosol.

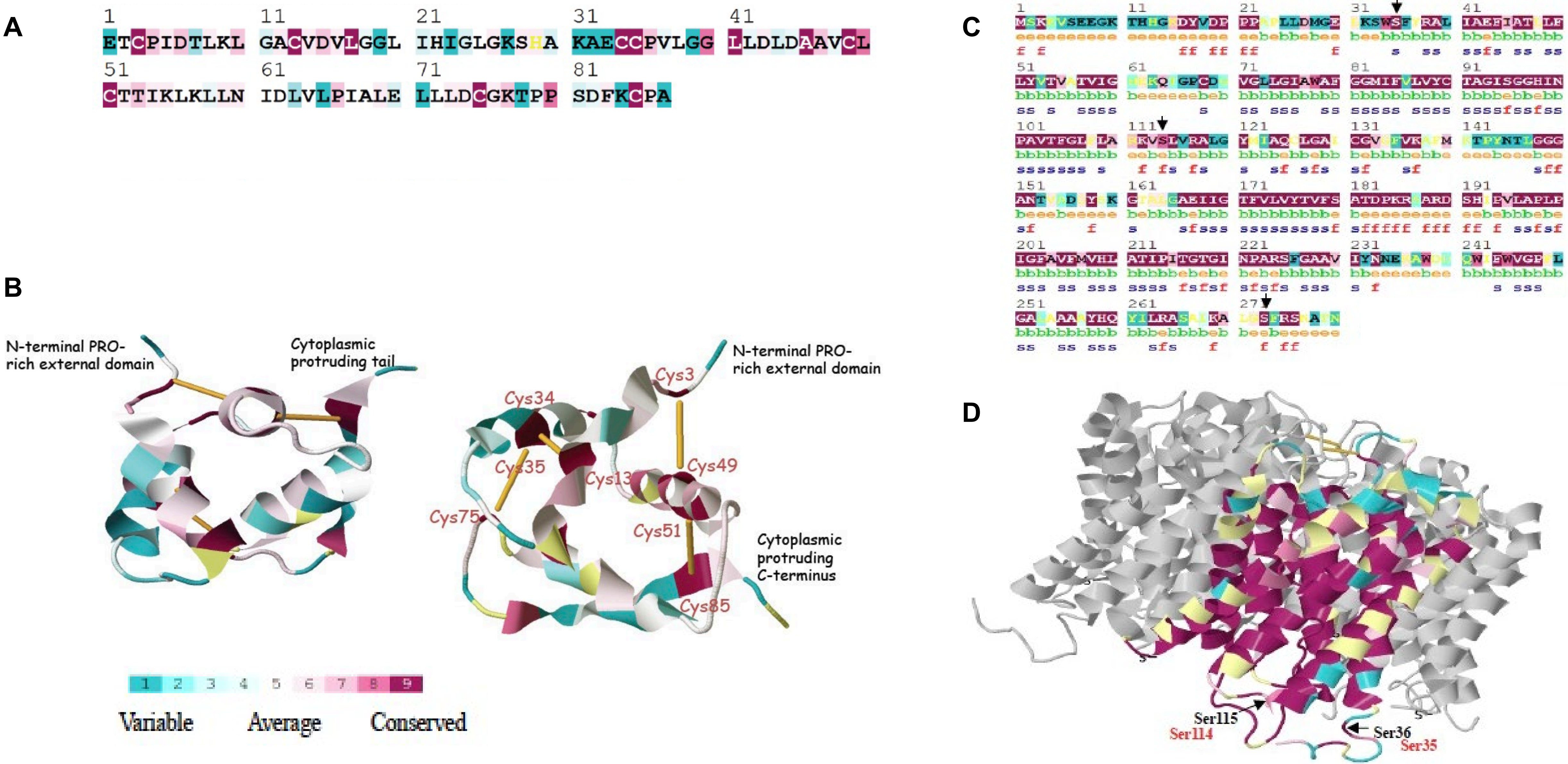


**Figure S2**
ConSurf analysis of evolutionary conservation of AtCWLP 8CM domain and AtPIP2;7.
(A) ConSurf analysis of amino acid sequence conservation based on a multiple alignment of the 8CM-domains of 29 Arabidopsis HyPRPs (including AtCWLP). At3g22120, At1g12090, At1g12100, At1g62500, At1g62510, At2g10940, At2g45180, At3g22142, At4g00165, At4g12470, At4g12480, At4g12490, At4g12500, At4g12510, At4g12520, At4g12530, At4g12545, At4g12550, At4g15160, At4g22460, At4g22470, At4g22485, At4g22490, At4g22513, At4g22517, At4g22520, At4g22160, At5g46890 and At5g46900) and four from other plants (maize B8QVC9, wheat Q4U1A2, soybean P24337, ragi P01087.2). The AtCWLP 8CM domain is displayed showing the Cys residues and 4 additional ones as the most evolutionary conserved residues.
(B) Predicted structure of AtCWLP transmembrane domain based on 1HYP 3D structure. Consurf analysis based on the crystal structure of GmHPS (PDB entry 1HYP, Baud et al. (1993) J Mol Biol 231, 877–887) sharing 36% identity with the 8CM domain of AtCWLP. The prediction shows a compact globular structure composed of four α-helices (left panel), stabilized by four disulfide bridges (yellow rods, right panel). AtCWLP Cys residues are numbered as indicated in the conservation plot (in A).
(C) Prediction of functionally and structurally important amino acid residues of AtPIP2;7 by the ConSurf server (Ashkenazy et al., 2010), based on Multiple Sequence Alignment (MSA) of 18 plant PIP2s (NCBI). The MSA included PIP2;7 (NP_195236.1), PIP2;8 (NP_179277.1), PIP2;6 (NP_181434.1), PIP2;5 (NP_191042.1), PIP2;2 (NP_181254.1), PIP2;3 (NP_181255.1) *of A. thaliana*; (XP_002869108.1), (XP_002884019.1) and (XP_002879778.1) of *A. lyrata*, (XP_002521292.1) and (XP_002530280.1) *of Ricinus cummunis*, (XP_002285157.1) of Vitis vinifera, (XP_002313510.1) of *Populus trichocarpa*, (XP_002461930.1) of *Sorghum bicolor*, (NP_001105026.1) of Zea mays, (NP_001059545.1) of *Oryza sativa*, (BAG68662.1) of *Tulipa gesneriana* and >gi|85544228|pdb|2B5F|D Chain D of spinach. The evolutionarily conserved regions and amino acid residues are colored according to the scale of conservation grades. Three highly-conserved Ser residues (S35, S114, and S273) of Arabidopsis aquaporins, corresponding to the tulip Ser residues (S35, S116, and S274) are marked by arrows (also in panel D).
(D) The ConSurf software was employed for obtaining a 3D model of AtPIP2;7, based on the crystal structure of spinach SoPIP2;1 in open conformation (PDB 2B5F). Amino acid residue numbering: black – spinach, red – Arabidopsis.

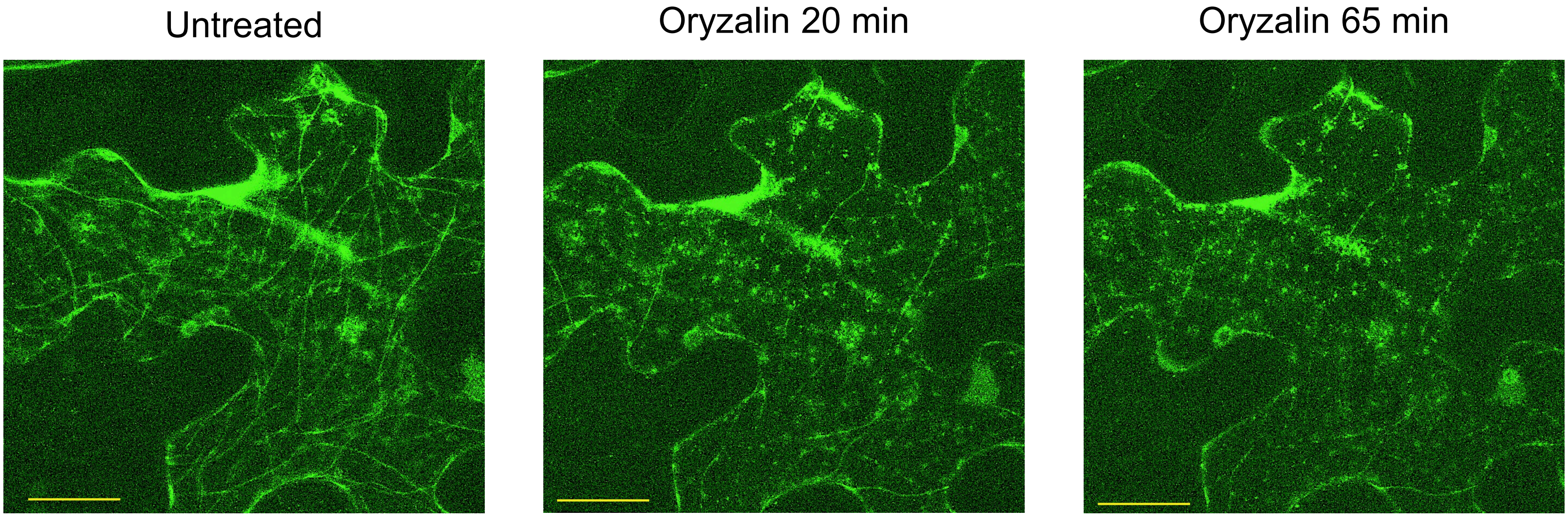


**Figure S3**
PRP940-GFP is not co-localized with microtubules. The effect of 5 µM oryzalin on PRP940 PM localization. PRP940-GFP was transiently expressed in *N. benthamiana* leaves. A detached piece of the injected area was studied under confocal microscope (LSM) prior and after the addition of 5 µM of microtubule disrupting agent oryzalin. The assembly of PRP940-GFP was not interrupted by oryzalin, while microtubules gradually disappeared. Scale bars = 20 µm.

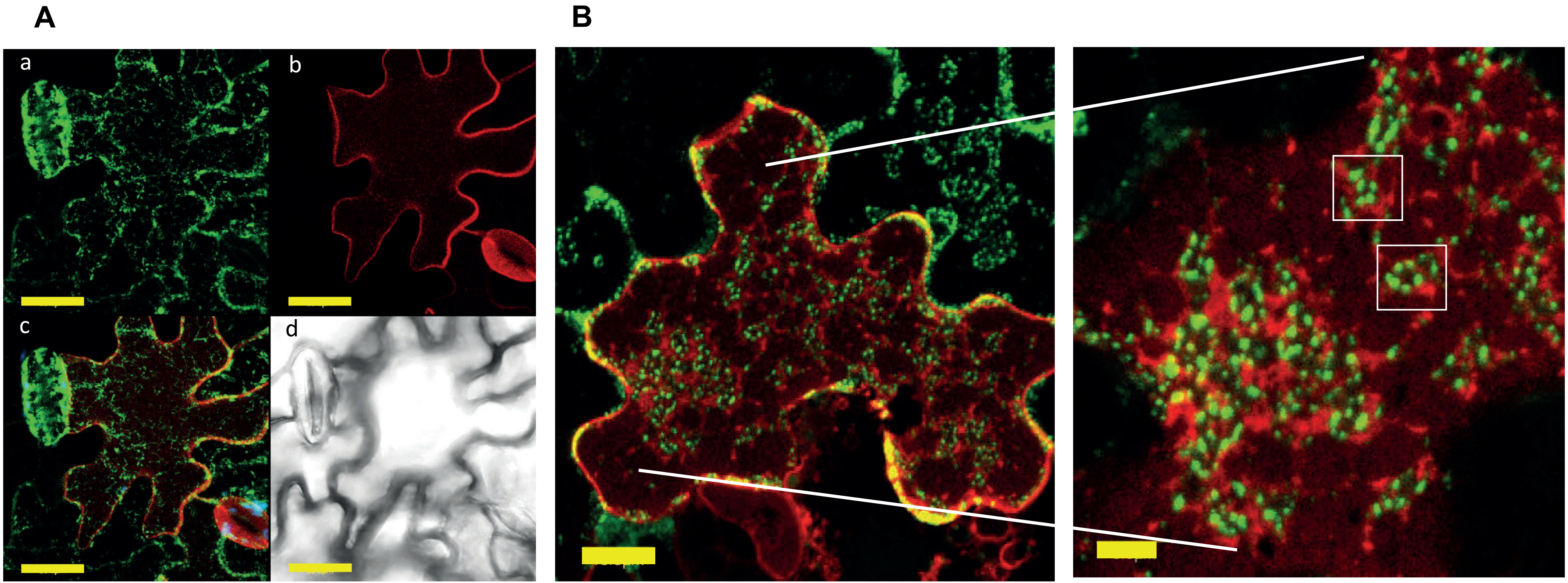


**Figure S4**
The effect of LatB on PM-localization patterns of CWLP-GFP and PIP2;1-mCherry ectopically expressed in transgenic Arabidopsis plants. (A) Untreated cells of abaxial leaf epidermis. (a) CWLP-GFP, (b) PIP2;1-mCherry, (c) merged, and (d) Nomarski images. Scale bars = 20 µm. (B) A similar cell after 60 min incubation with 25 µM LatB. Scale bar = 10 µm. The right enlarged image shows the formation of large endocytic vesicle-like structures containing CWLP-GFP clusters (indicated in white frames) or PIP2;1-mCherry or both, resulting from the LatB treatment. Scale bars = 5 µm.

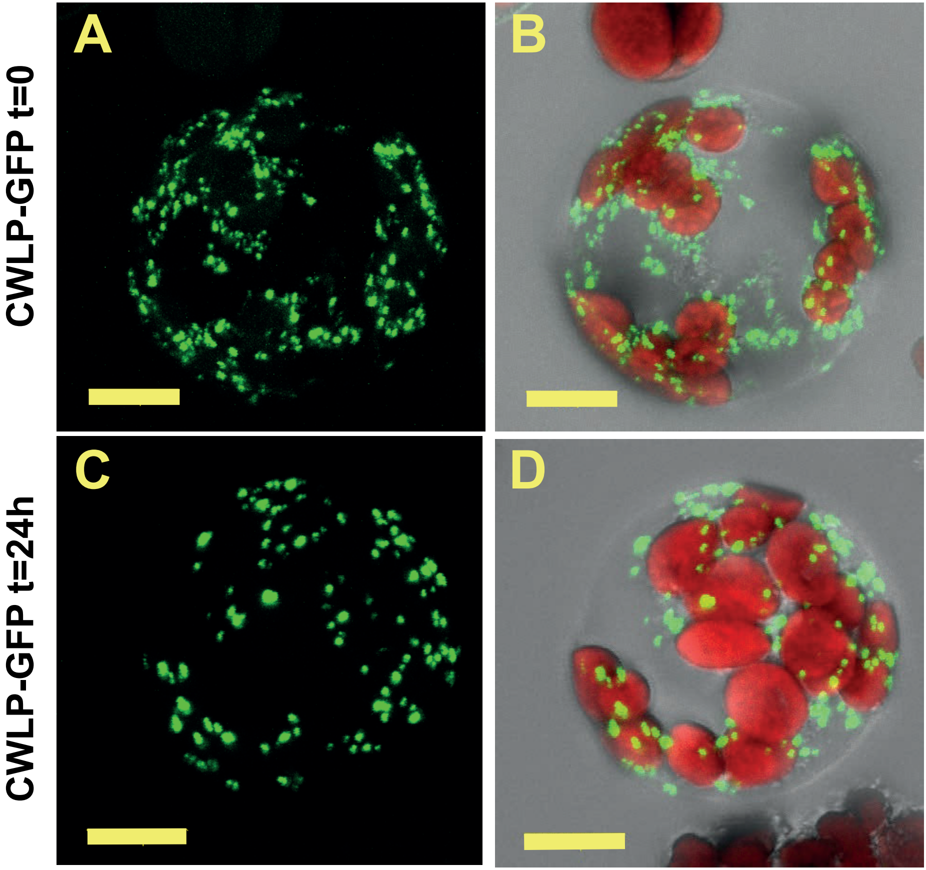


**Figure S5**

Clustering of CWLP-GFP in the PM is not affected in CW-free protoplasts prepared from leaves of CWLP-GFP expressing Arabidopsis plants. (A) and (B) Protoplasts after removing the cell wall by digesting enzymes. (C) and (D): CWLP-GFP clustering in the PM 24 h after protoplast preparation. GFP and chloroplast fluorescence is shown in green and red, respectively. Bars = 10 µm.

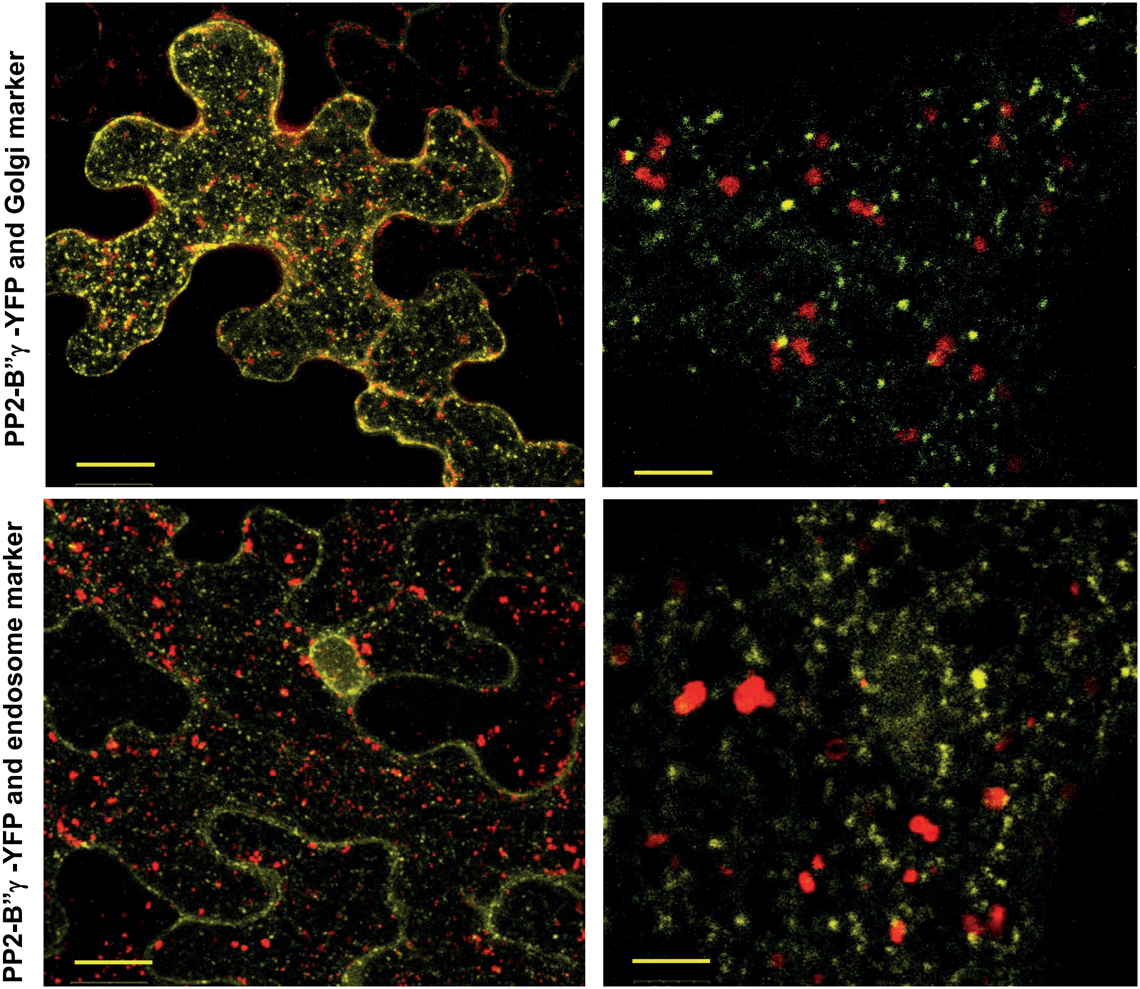


**Figure S6**
PP2A-B”γ-YFP is not co-localized with Golgi and endosomal markers upon transient expression in *N. benthamiana* leaves. Upper panel: Co-expression of PP2A-B”γ-YFP and the Golgi marker GmMannosidase1-mCherry. Lower panel: Co-expression of PP2A-B”γ-YFP and the endosomal marker FYVE-DsRed. Left image bars 20 μm, and right image bars 5 μm.

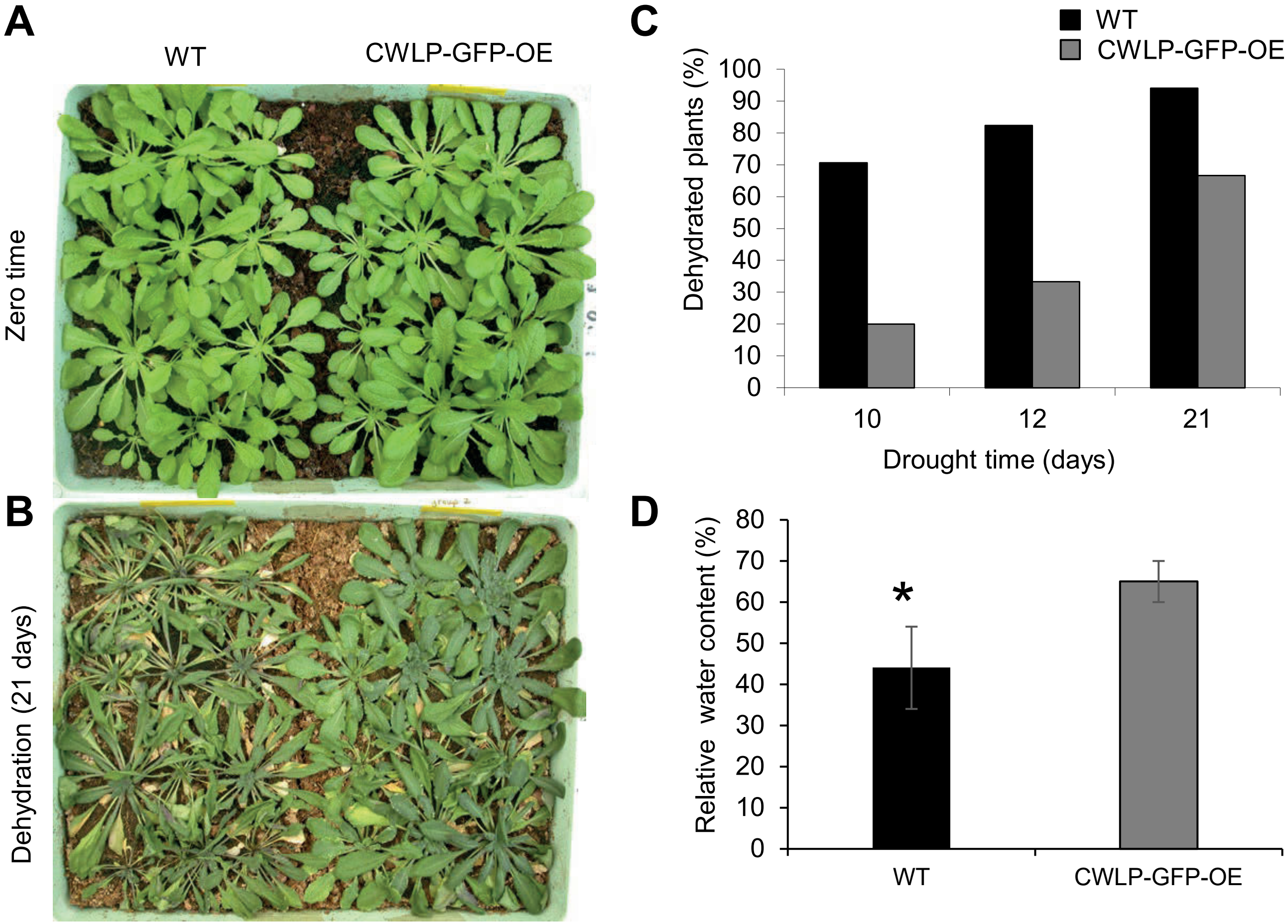


**Figure S7**
CWLP overexpression increases tolerance to water shortage. Three-month-old WT and CWLP-GFP-OE plants were exposed to dehydration by withholding irrigation. (A) WT (left) and CWLP-GFP-OE (right) plants after the last irrigation (zero time point). (B) WT (left) and CWLP-GFP-OE (right) plants after 21 days of water withholding. (C) Percentage of dehydrated plants, corresponding to the experiment shown in B. WT - Black columns, CWLP-GFP-OE - Grey columns (D) Relative water content (RWC) of WT and CWLP-GFP-OE rosette leaves after 20 days of dehydration. The values represent the average of 15 leaves (3 leaves taken from each of 5 different plants). The experiment was repeated three times.

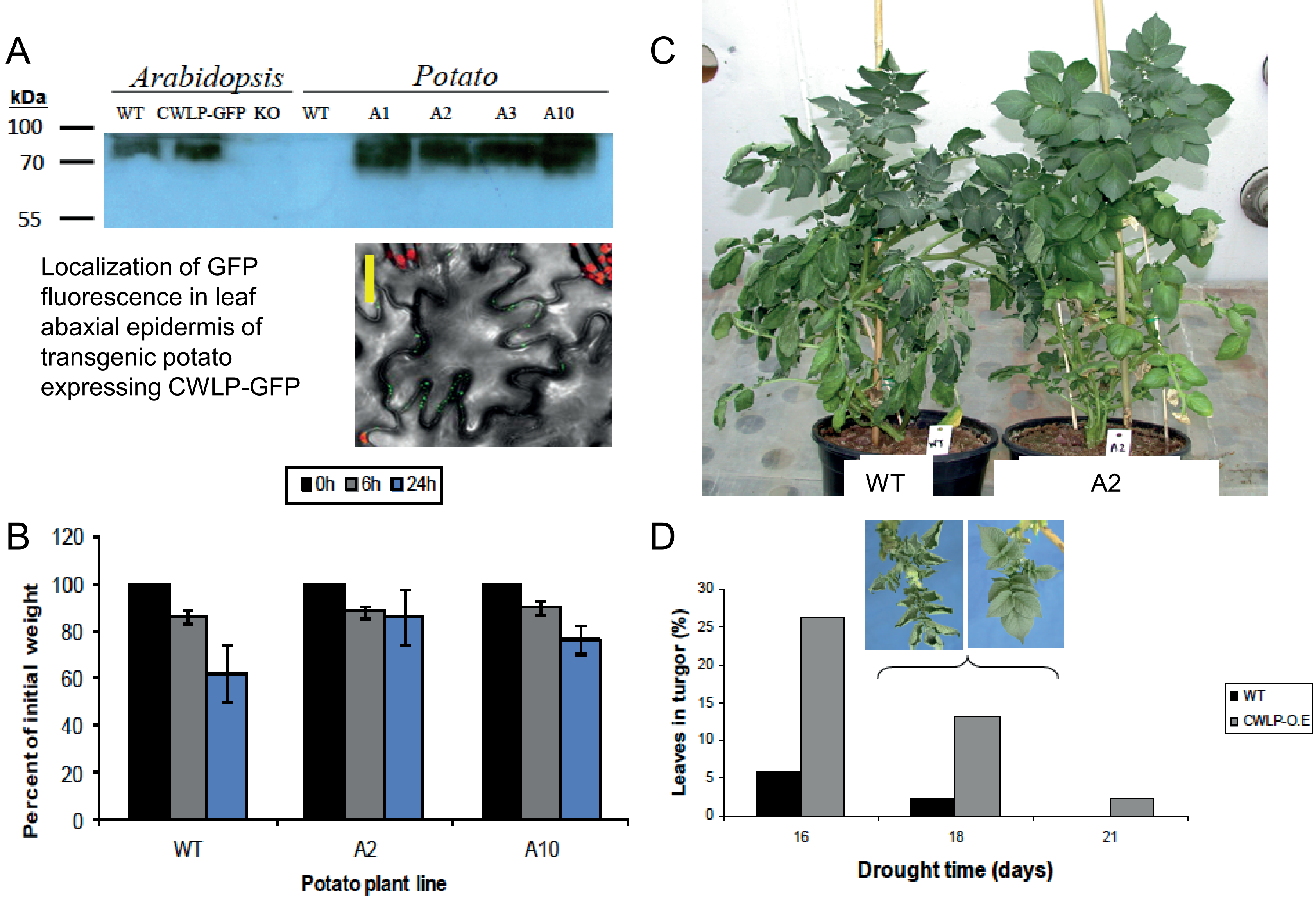


**Figure S8**

Overexpression of CWLP-GFP in potato confers improved dehydration tolerance.

(A) CWLP-GFP expression in Arabidopsis and potato analyzed by western blotting with polyclonal anti-CWLP antibody. WT (wild type Col-0) or potato (Desiree) plants, CWLP-GFP – Arabidopsis plants overexpressing CWLP-GFP, KO – *cwlp*, GABI_382D01 T-DNA K/O mutant, A1 to A3 and A10 – potato plants overexpressing CWLP-GFP (CWLP-O.E). Scale bar = 20 µm.

(B) Water loss during 24 days of water withholding in potato plants. Water loss is lower in CWLP-O.E compared with WT plants.

(C) Comparison of WT and A2 (CWLP-O.E) plants after 16 days of water withholding.

(D) Turgor loss kinetics; comparison of WT and potato A2 plants. The insert magnifies leaves of WT (left) and A2 (right) plants on the 18th day of water withholding.

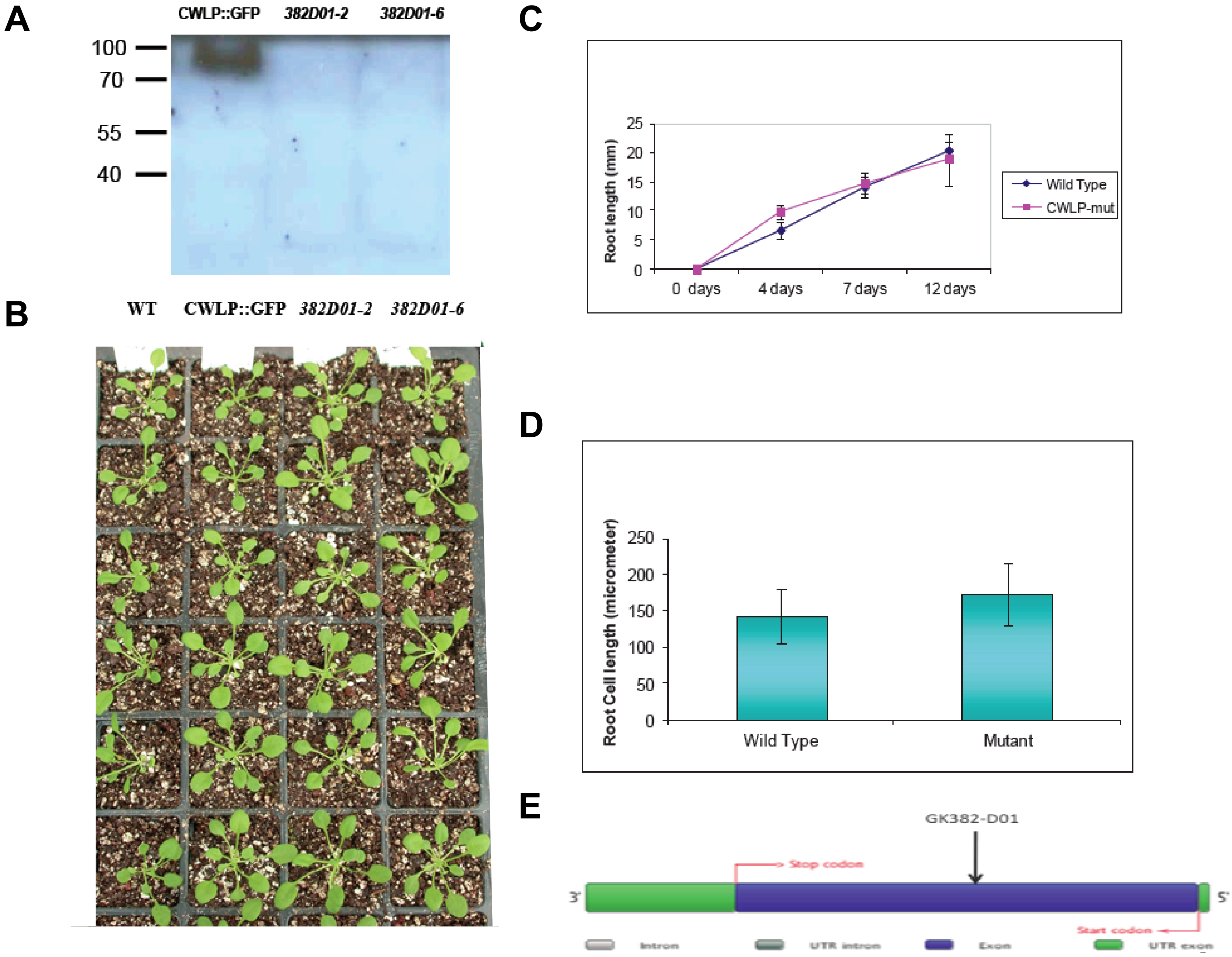


**Figure S9**
The phenotype of *cwlp* T-DNA knockout line does not differ from that of WT. (A) No CWLP protein is detectable in *cwlp* (GABI_382D01) homozygous plants (two independent F3 offspring 382D01-2 and 382D01-6). Western blot probed with polyclonal anti-CWLP antibody. (B) No morphological difference in growth of WT and *cwlp* seedlings. (C) No significant difference in root growth of WT, CWLP-GFP and *cwlp* seedlings recorded during 12 days after germination (DAG). (D) No significant difference in cell length in the root elongation zone between the WT and the *cwlp* plants. Measurements performed by confocal microscope. (E) Schematic map of T-DNA insertion in the *CWLP* gene in the GK382-D01 knockout mutant.

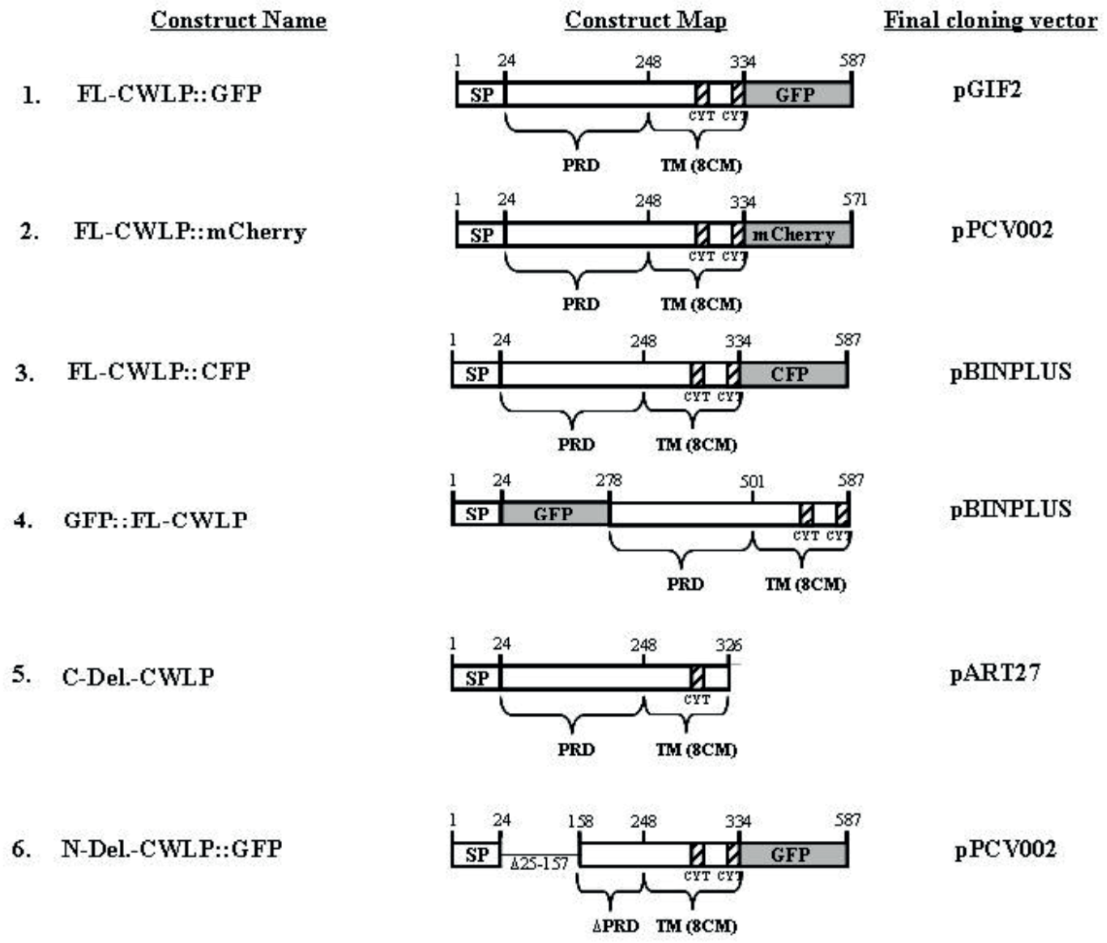


**Figure S10**
AtCWLP constructs used in this study. For additional information see Supplemental Information.
